# Soluble SORL1 in cerebrospinal fluid as a marker for functional impact of rare SORL1 variants

**DOI:** 10.1101/2025.08.05.668459

**Authors:** Matthijs W. J. de Waal, Sven J. van der Lee, Melanie Lunding, Lynn Boonkamp, Nolan Barrett, Giulia Monti, Anne Mette G. Jensen, Christian B. Vaegter, Jan Raska, Sona Cesnarikova, Jiri Sedmik, Calvin Trieu, Marjan M. Weiss, Rosalina van Spaendonk, Lisa Vermunt, Georgii Ozhegov, Niccolo Tesi, Marc Hulsman, Dovile Januliene, Arne Moller, Dasa Bohaciakova, Wiesje M. van der Flier, Olav M. Andersen, Charlotte E. Teunissen, Henne Holstege

## Abstract

**Background:** As part of the retromer, sortilin-related receptor (SORL1) sorts cargo proteins away from the endosome to the trans-Golgi network or to the cell surface, where SORL1 is cleaved into sSORL1 and shed into the interstitial fluid and CSF. SORL1’s cargo includes APP and Aβ, which may explain why protein-truncating genetic variants (PTVs) in *SORL1* are observed almost exclusively in Alzheimer’s Disease (AD) patients, and that rare, predicted pathogenic missense variants have been associated with a 10-fold increased risk of AD. However, functional evidence supporting variant pathogenicity is warranted. Here we investigated whether soluble SORL1 (sSORL1) concentrations in cerebrospinal fluid (CSF) offer a potential in vivo biomarker to support impaired SORL1 function in variant carriers.

**Methods:** Using an ELISA assay for SORL1 (ABCAM), we determined sSORL1 concentrations in CSF from 218 participants of the Alzheimer Dementia Cohort (ADC) (54% females). We compared sSORL1 in CSF derived from 90 carriers of diverse *SORL1* variants with concentrations observed in 78 *SORL1*-WT AD patients, and 50 *SORL1*-WT controls for whom CSF-pTau-181, CSF-tTau, CSF-Aβ42 concentrations were available. In a subset of 36 individuals, we used Western blotting (WB) to validate sSORL1 concentrations as determined by ELISA.

**Results:** CSF-sSORL1 concentrations did not differ between *SORL1*-WT AD patients and controls. While CSF-sSORL1 did not correlate with CSF-Aβ42 concentrations in *SORL1*-WT AD patients (p=0.62), it correlated with sCSF-ptau-181 (p=9.7×10^−6^). The mean CSF-sSORL1 concentration in the SORL1 WT AD cases and controls was 466 pg/ml, with wide variance (SD = 133). Comparatively, concentrations were significantly lower in PTV carriers (260 pg/ml, p=3.2×10^−7^) and in carriers of predicted damaging *SORL1* missense variants (323 pg/ml, p=2.4×10^−7^). CSF-sSORL1 concentrations measured by ELISA correlated strongly with concentrations estimated by WB (ρ=0.552; p=5.0×10^−4^).

**Conclusion:** CSF-sSORL1 increases with CSF-ptau, suggesting that increased SORL1-retromer activity may serve to rescue cellular stress associated with AD-related processes. However, impairing *SORL1* genetic variants may preclude SORL1 trafficking to the cell surface, as supported by lower CSF-sSORL1 protein concentrations in carriers. While further refinements are necessary, we present first evidence for the applicability of ELISA-based quantification of sSORL1 in CSF to evaluate the functional impact of rare *SORL1* variants.

## Introduction

Sortilin-related receptor (SORL1 also known as SORLA and LR11) operates in the endolysosomal pathway as a sorting receptor of various proteins, with a main function to traffic cargo proteins from the endosome to the Golgi or for recycling of cargo from the endosome back to the cell surface (1, 2). Genetic variants that impair SORL1 function have been linked with an increased risk of developing Alzheimer’s disease (AD)(3-5). In the context of AD, SORL1 can regulate the proteolysis of amyloid precursor protein (APP) into Amyloid-Beta (Aβ) by controlling the transport of APP to the endosome (6-10). In addition, SORL1 facilitates the trafficking of cargo-Aβ to lysosome for degradation (11). *SORL1* protein-truncating variants (PTVs) have been observed almost exclusively in AD cases, supporting *SORL1* haploinsufficiency (3, 12). However, the majority of *SORL1* variants are rare missense variants, with diverse effects on SORL1 function (13, 14). While most variants are likely benign, a subset of variants have strong risk-increasing effects on AD (12).

Determining of the pathogenicity of *SORL1* variants is challenging, as variants are rare and pedigrees of *SORL1* variant carriers are commonly small (15-18). We recently proposed a prioritization scheme of *SORL1* missense variants according to a ‘disease mutation domain mapping’ approach (DMDM), a comprehensive manual effort, taking in-depth knowledge of SORL1 functional domains into account(19). Using this approach, we leveraged mutations in proteins associated with monogenic diseases that share homologous domains with SORL1 to identify domain positions where mutations are predicted to be pathogenic. This led to the categorization of missense variants into high-moderate, low- and no-priority missense variants. Testing the effect on AD risk in a large sequencing dataset from AD cases and controls (12) indicated that high-priority variants associated with a 10-fold increased risk of early onset AD and a 6-fold increased risk of overall AD. In contrast, moderate-, low- and no-priority missense variants did not associate with AD risk. In addition to DMDM, variant prioritization may further benefit from a functional readout to support DMDM prediction of pathogenicity. Here, we leveraged the observation that functional SORL1 traffics to the cell-surface, where the ectodomain of the receptor is cleaved by ADAM17 (TACE) into extracellular sSORL1, which is released in the interstitial space and CSF (20). Impaired trafficking of the SORL1 receptor might be reflected in lower levels of SORL1 that reach the cell surface, and thus lower levels of cleaved sSORL1 as recently shown using WB (21). Therefore, we explored whether ELISA for CSF sSORL1 can serve as an effective quantitative *in vivo* biomarker for pathogenic *SORL1* missense mutations. We first analytically validated a commercially available immunoassay to measure sSORL1 in CSF. Next, we measured sSORL1 concentrations in human CSF samples from AD patients who carried *SORL1* genetic variants with varying predicted pathogenicity, as well as from non-variant carriers. Finally, we compared these CSF concentrations with those measured using Western blotting to assess the consistency in our findings.

## Methods

### Study cohort

For this study, we selected 218 participants from the Amsterdam Dementia Cohort (ADC), which includes patients who visited the memory clinic at the Alzheimer Center, Amsterdam University Medical Center, The Netherlands, and underwent a diagnostic work-up (22). We selected participants with available cerebrospinal fluid (CSF) samples and genetic information and divided them into three groups (**see Fig. 1**). The first group consisted of 90 participants carrying rare genetic variants in the *SORL1* gene. Most participants within this group were diagnosed with dementia due to AD (n=71), while others had diagnoses of mild cognitive impairment (MCI) (n=9), vascular dementia (n=2), psychiatric disorder (n=2), SCD (n=5), non-neurodegenerative neurological disease (n=1). Clinical diagnosis was made through consensus-based, multidisciplinary meeting. AD-type dementia diagnosis was according to the clinical criteria formulated by the National Institute of Neurological and Communicative Disorders and Stroke—Alzheimer’s Disease and Related Disorders Association (NINCDS-ADRDA) and based on National Institute of Aging–Alzheimer association (NIA-AA) (23, 24). AD diagnoses were supported by biomarker confirmation of amyloid-positive status (25), determined through CSF analysis (n=55) or PET imaging (n=15), except for one carrier whose amyloid status could not be confirmed. MCI diagnosis was based on international consensus criteria (26) and vascular dementia was diagnosed according to the NINDS-AIREN criteria (27). Diagnoses for psychiatric or non-neurodegenerative neurological disorders were based on the diagnostic work-up. The second group consisted of *78 SORL1* wild-type AD patients (65 CSF and 13 PET confirmed amyloid-positive) that were selected to match optimally with the 90 carriers by age and sex. The third group consisted of 50 randomly selected cognitively normal *SORL1* wild-type controls, all of whom showed no signs of cognitive impairment during their diagnostic work-up. Cognitive impairment was assessed using the Mini-Mental State Examination (MMSE)(28), with data available for all patients. In addition, APOE genotype information was available for all patients. All patients gave informed consent for the use of their medical data and biomaterial. The Medical Ethics Committee of the Amsterdam UMC approved this study in accordance with the declaration of Helsinki.

**Fig 1.**
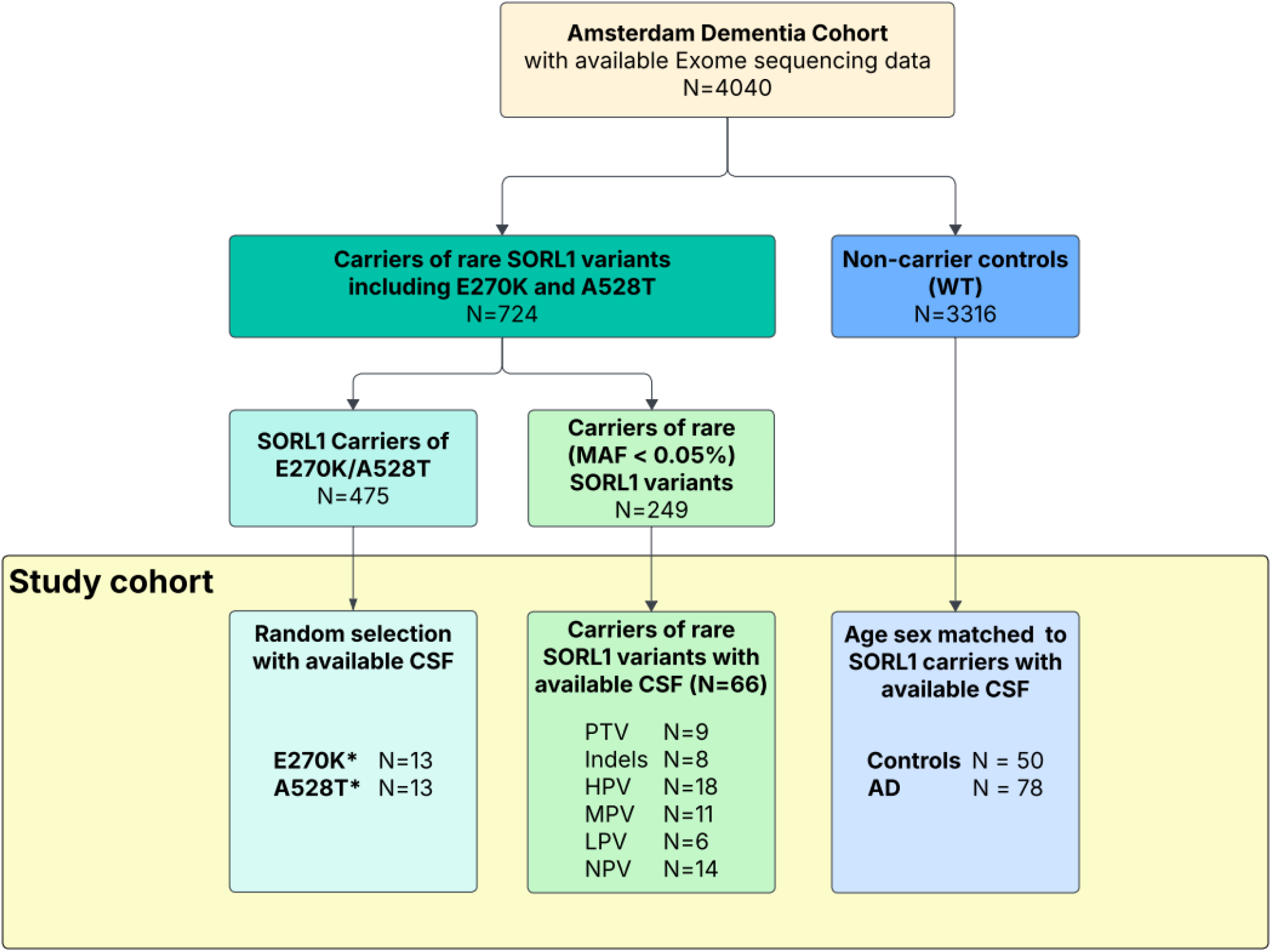
Selection of the study cohort. Participants were included from the Amsterdam Dementia Cohort. *SORL1* variant carrier status was determined using whole-exome sequencing. Samples for which CSF was available were included in the Study cohort, WT-SORL1 included controls and AD patients (amyloid-confirmed). Abbreviations: A528T = p.Ala528Thr; CSF = Cerebrospinal Fluid; E270K = p.Glu270Lys; HPV = High priority variants; Indels = in-frame insertion/deletion; LPV = Low priority variant; MAF = Minor allele frequency; MPV = moderate priority variant; NPV = No priority variant; PTV = Protein truncating variant; WT = Wild type. *2 carriers both carried an E270K and A528T mutation and were included in both groups.

### *SORL1* genetic variant detection

*SORL1* variants were determined by whole exome sequencing (WES)(4, 5) and stratified into different pathogenicity categories according to previously described selection criteria (12, 19). Protein truncating variants (PTVs), included in-frame insertion/deletion variants (in-frame indels) and nonsense variants. Missense variants were stratified according to previously described Domain Mapping of Disease Mutations approach (DMDM) (12, 29), into high priority missense variants (HPV), moderate priority missense variants (MPV), low priority missense variants (LPV) and no priority missense variants (NPV). Since no clear disease-associated mutations have been found in any of the VPS10p family members, it was not possible to perform DMDM for the VPS10p/10CC domain, such that HPVs in this domain were based on having a REVEL score >0.5 and MAF (>0.05%; gnomAD)(30, 31).

### CSF sample collection and measures

CSF samples were obtained as part of routine clinical care by lumbar puncture using a 20/25-gauge needle and syringe between the L3/L4, L4/L5, or L5/S1 intervertebral space, collected in polypropylene tubes, and processed as previously described (32). Concentrations of the CSF AD core biomarkers (Aβ42 (n=216), phosphorylated Tau-181 (pTau-181; n=162) and total Tau (tTau; N=162)) were measured in the Amsterdam UMC using commercial kits: Elecsys Aβ42, pTau (181P) and tTAU CSF assays (Roche Diagnostics, Basel, Switzerland) or INNOTEST β-Amyloid(1-42), pTau (181P) and hTAU Ag (Fujirebio, Gent, Belgium). Aβ42 concentrations measured on INNOTEST were corrected for a drifting effect in biomarker concentrations that appeared throughout the years (33). Abnormal amyloid status (A+) was determined based on previously defined abnormal assay-specific concentration cut-offs in cerebrospinal fluid (CSF): for INNOTEST, an Aβ42 level of <813 pg/ml was considered abnormal, while for Elecsys, an a pTau-181/Aβ42 ratio of ≥0.02 was considered abnormal (33, 34).

### Validation and application of SORL1 ELISA in CSF

The quantification of sSORL1 was performed with a commercially available ELISA kit (ab282864, ABCAM, Cambridge, UK) according to the manufacturer’s protocol, which was validated for Heparin Plasma, Cell Culture Supernatant, Serum, Cell Lysate, and EDTA Plasma. Here, we analytically validated this assay for 10x diluted CSF, following standard protocols (35). We evaluated the assays’ precision, parallelism, dilution linearity, recovery, specificity, target sensitivity and sample stability. Details of the different parameters can be found in the **Supplemental Methods**. This includes information on cell culture and SORL1 constructs used for specificity. Absorbance was measured with the EPOCH2NS reader (BioTek, Agilent, Santa Clara, US) using Gen5 reader software (version 3.12). After validation, we applied the assay to 218 CSF samples as analyzed here.

### Western Blot (WB) analysis

For 37 *SORL1* variant carriers, we compared the ELISA results with the relative sSORL1 CSF concentrations (rel-sSORL1) determined by WB. We determined the sSORL1 CSF concentrations in the variant carriers relative to concentrations of optimally matched with *SORL1* WT AD patients, based on gender, APOE genotype and age at AD onset (**Table S1**). Details of the estimation of the rel-sSORL1 concentration and generation of sSORL1 standard protein are described in the **Supplemental Methods**. In short, equal volumes of CSF or sSORL1 standard protein were separated by SDS-PAGE using NuPAGE 4-12% Bis Tris gels (NP0321BOX, Thermo Fischer, Waltham, US) and then transferred to nitrocellulose membranes (Amersham GE Healthcase Life Sciences, Chicago, US). Unspecific binding was blocked (0.25 M Tris-Base, 2.5 M NaCl, 2% skimmed milk powder, 2% Tween-20) for 1 h at room temperature, and incubated overnight at 4°C with mouse anti-LR11 primary antibody (611860, BD Biosciences, Franklin Lakes, US). The following day, the membranes were incubated in polyclonal rabbit anti-mouse HRP secondary antibody (P0260, 1:1500; Agilent Dako) for 1 h at room temperature. After washing membranes in washing buffer (0.2 mM CaCl_2_, 0.1 mM MgCl_2_, 1 mM HEPES, 14 mM NaCl, 0.2% skimmed milk powder, 0.05% Tween 20) for a total of 25 min, proteins were detected with SuperSignal West Femto Maximum Sensitivity Substrate (34094, Thermo Fisher), and images were captured using LAS-4000 chemiluminiscent imager (GE Healthcare). Signals were quantified with Multi Gauge software (Fujifilm Life Sciences, Tokyo, Japan).

### Statistical analysis

We analyzed the association between CSF-sSORL1 concentrations and pathogenicity or *SORL1* variants in two ways. First, we used robust linear regression to investigate the relationship between CSF-sSORL1 concentrations and variant-effect on AD risk, as estimated for each of eight variant pathogenicity categories, while adjusting for age and sex. Rare missense variants were assigned to the (1) high-, (2) moderate-, (3) low- and (4) no-priority group, as estimated by DMDM in our previous study (12). Truncating variants were assigned to the (5) PTV group and individuals without *SORL1* variants were assigned to the (6) wild-type (WT) group. The risk level associated with each pathogenicity category was the natural logarithm of odds ratios associated with it, as estimated by our previous study (12).

Second, we performed binary robust linear regression to compare CSF-sSORL1 concentrations across each pathogenicity category with those in the *SORL1* WT control group (n = 128), as well as with the AD WT group (n = 78) and cognitively unimpaired controls (n = 50). Statistical significance was set at a Bonferroni-corrected threshold of 0.00625 (0.05/8 groups).

Then we associated sSORL1 concentrations with (1) age (adjusted for sex), (2) sex (adjusted for age), (3) *APOE* genotypes, (4) MMSE, and (5) AB42 and (6) tTau and (7) pTau-181, using robust linear regressions adjusting for age and sex. The threshold for statistical significance was set at 0.00714, to correct for multiple testing (0.05/7 tests). In this analysis, we considered only SORL1 WT AD cases, and excluded controls and *SORL1* mutation carriers, to minimize potential confounding effect from *SORL1*-related deficiencies on AD pathology. Potential interactions of *APOE ε4* genotype on sSORL1 concentrations across carrier groups were calculated incorporating the interaction term between *APOE ε4* and the *SORL1* genotype groups (sSORL1 concentration ~ pathogenicity category group ^*^ *APOE ε4* genotype).

We aimed to define the lower end of the normal range of sSORL1 concentrations. Values below this threshold should be enriched for pathogenic variant carriers. For this we estimated the 1st percentile of sSORL1 concentrations (pg/ml) in the control group and its 95% confidence interval (CI). We performed a bootstrapping procedure with 1,000 resampling iterations. In each iteration, we drew a new dataset by randomly sampling (with replacement) from the observed sSORL1 values in the control group, maintaining the original sample size. The 1st percentile was calculated for each bootstrap sample, and the 95% CI was determined as the 2.5th and 97.5th percentiles of the bootstrapped distribution. Relative sSORL1 concentrations in CSF quantified via Western Blot (WB), were compared against a reference median value of 100, representing the control group using Wilcoxon signed-rank test.

## Results

### SORL1 ELISA analytical validation

We tested the performance of the sSORL1 ELISA kit for CSF-samples, a matrix that was not included in the specifications of the manufacturer. The lower limit of detection (LLOD) for this assay is 55 pg/ml. Precision of the assay in CSF was robust, with a 3.9% intra-assay variability (low QC = 4.4%, medium QC = 3.4% and high QC = 3.8 and a 10.0% inter-assay variability (low QC = 12.0%, medium QC = 9.8% and high QC = 8.2%). Parallelism (**Fig. 2A**), linearity (**Fig. 2B**), recovery (**Fig. 2C**) and freeze-thaw stability (**fig. 2D**) were all within the 85-115% acceptable range. Thus, the assay allows the estimation of variation in CSF-sSORL1 concentrations in CSF.

**Fig 2.**
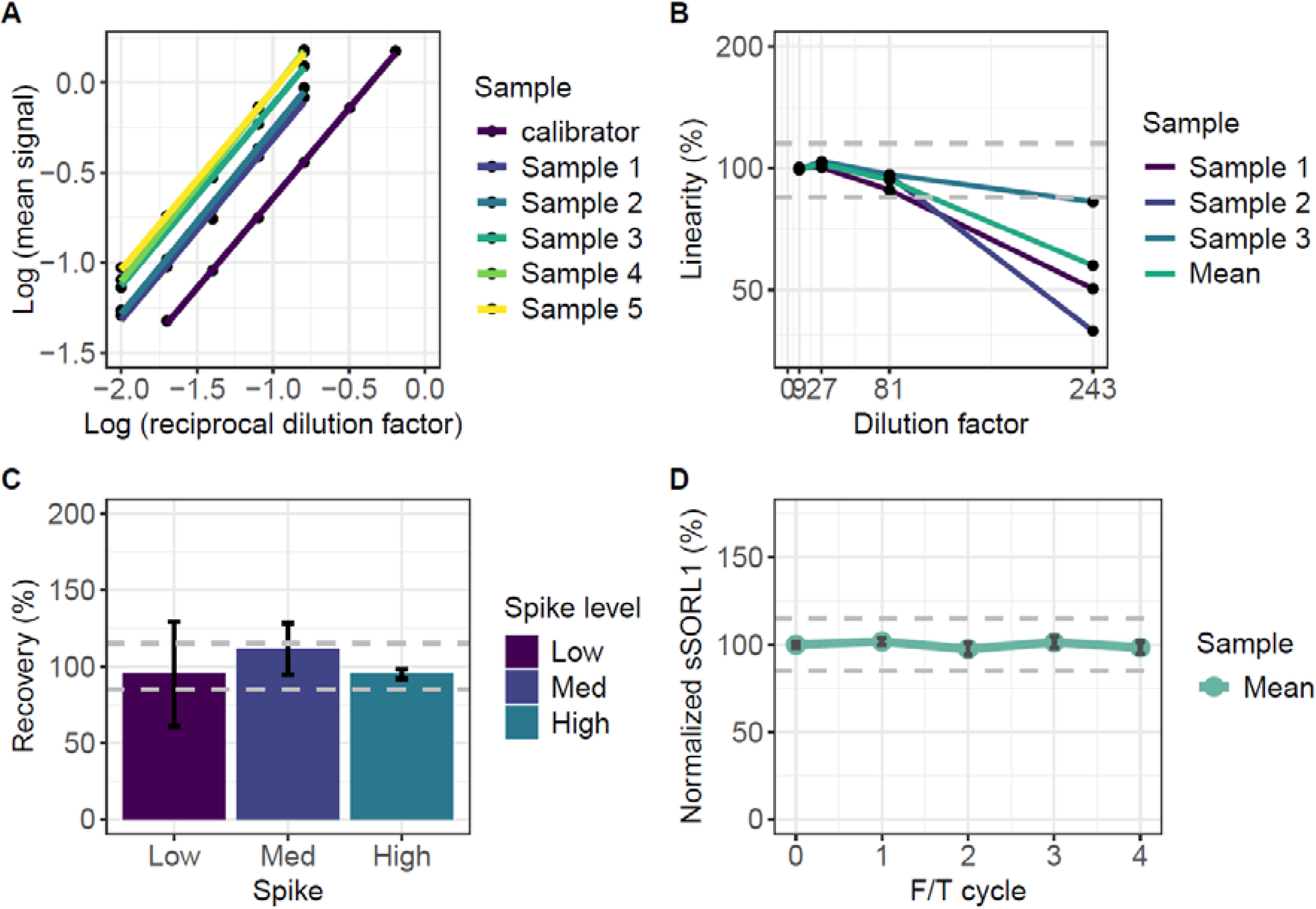
Technical validation of the CSF-sSORL1 ELISA assay. A.) Parallelism indicates that 2-fold serial dilution of sSORL1 measurements are parallel to the measurement of the standard curve. B.) Dilution linearity indicates that samples remain reliably measurable within a range of 9-fold to 81-fold dilution. C.) The recovery of low, medium, and high spike concentrations falls within the expected range for accurate measurement. D.) Sample stability demonstrates that CSF samples subjected to 4 freeze/thaw cycles still yield similar concentrations.

Then, we examined specificity/cross-reactivity. According to the manufacturer, the antibodies for the ELISA assay were raised against a SORL1 recombinant fragment containing amino acid residue 82 to 755 of the SORL1 protein, which represents the VPS10p domain. This domain characterizes the VPS10p family of proteins (SORL1, SORT1, SORCS1, SORCS2 and SORCS3). To test for cross-reactivity of the ELISA with other VPS10p family members, we spiked solutions of recombinant SORL1 protein with defined concentrations of recombinant SORT1 or SORCS2 (see **Supplemental Methods**). We did not observe an increase in the sSORL1 concentration after spiking (**Fig. S1**), suggesting that there was no cross-reactivity. In addition, the assay was able to detect different full size SORL1 recombinant proteins, including one produced in-house (residue 82-2135) and another commercially available (residue 82-2135) (**Fig. S2**); see details in **Supplemental Methods**. Finally, to determine potential nonspecific signal, we measured the media of different *SORL1* KO cell lines, which should be devoid of any sSORL1 protein. Indeed, we observed almost no signal for the KO condition, while the WT and the mock cell lines showed good expression (**Fig. S3**). Overall, these results demonstrate the robustness and specificity of the assay for detecting sSORL1, without cross-reactivity to soluble cleaved products of other VPS10p domain-containing proteins.

### High-Risk *SORL1* Missense Variants Exhibit Lower CSF-sSORL1 Levels Similar to PTVs

To assess whether *SORL1* variant carriers had different sSORL1 concentrations compared to WT-*SORL1* carriers, we measured sSORLA concentrations in CSF derived from (1) 50 non-demented controls (average age at sample collection is 61 (SD=6), 38% female); (2) 78 *SORL1*-WT AD patients (average age at sample collection is 63, 66% female); (3) 90 *SORL1* variant carriers (average age at sample collection is 64 (SD=7), 50% female). See **Table 1** for demographics.

**Table 1.**
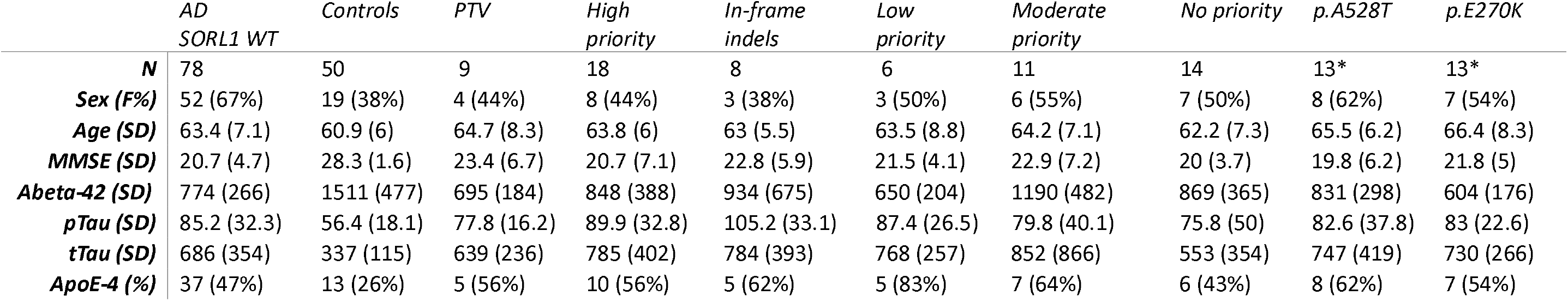
demographics of the cohort. *2 carriers have both an A528T variant and a E270K variant and are therefore included in both groups. AD = Alzheimer’s Disease, PTV = Protein truncating variant, In-frame indels = in-frame insertion deletion variant, MMSE = Mini-mental state examination, F = Female, SD = standard deviation.

|  | *AD*<br>*SORL1 WT* | *Controls* | *PTV* | *High*<br>*priority* | *In-frame*<br>*indels* | *Low*<br>*priority* | *Moderate*<br>*priority* | *No priority* | *p.A528T* | *p.E270K* |
| --- | --- | --- | --- | --- | --- | --- | --- | --- | --- | --- |
| ***N*** | 78 | 50 | 9 | 18 | 8 | 6 | 11 | 14 | 13* | 13* |
| ***Sex (F%)*** | 52 (67%) | 19 (38%) | 4 (44%) | 8 (44%) | 3 (38%) | 3 (50%) | 6 (55%) | 7 (50%) | 8 (62%) | 7 (54%) |
| ***Age (SD)*** | 63.4 (7.1) | 60.9 (6) | 64.7 (8.3) | 63.8 (6) | 63 (5.5) | 63.5 (8.8) | 64.2 (7.1) | 62.2 (7.3) | 65.5 (6.2) | 66.4 (8.3) |
| ***MMSE (SD)*** | 20.7 (4.7) | 28.3 (1.6) | 23.4 (6.7) | 20.7 (7.1) | 22.8 (5.9) | 21.5 (4.1) | 22.9 (7.2) | 20 (3.7) | 19.8 (6.2) | 21.8 (5) |
| ***Abeta-42 (SD)*** | 774 (266) | 1511 (477) | 695 (184) | 848 (388) | 934 (675) | 650 (204) | 1190 (482) | 869 (365) | 831 (298) | 604 (176) |
| ***pTau (SD)*** | 85.2 (32.3) | 56.4 (18.1) | 77.8 (16.2) | 89.9 (32.8) | 105.2 (33.1) | 87.4 (26.5) | 79.8 (40.1) | 75.8 (50) | 82.6 (37.8) | 83 (22.6) |
| ***tTau (SD)*** | 686 (354) | 337 (115) | 639 (236) | 785 (402) | 784 (393) | 768 (257) | 852 (866) | 553 (354) | 747 (419) | 730 (266) |
| ***ApoE-4 (%)*** | 37 (47%) | 13 (26%) | 5 (56%) | 10 (56%) | 5 (62%) | 5 (83%) | 7 (64%) | 6 (43%) | 8 (62%) | 7 (54%) |

We categorized the CSF samples from the 90 *SORL1* variant carriers according to predicted variant effects (listed in **Table S1**): 9 protein truncating variants (PTV)s, 18 rare high priority variants (HPVs), 11 rare moderate priority variants (MPVs), 6 rare low priority variants (LPVs), rare 14 no-priority variants (NPVs), rare 8 in-frame indels, 11 carriers of the common E270K variant (rs117260922), 11 carriers of the common A528T variant (rs2298813), and 2 individuals who carried both the E270K and A528T common variants. The remaining 128 CSF samples were derived from individuals who were WT for *SORL1*. CSF sSORL1 concentrations varied widely among individuals (**Fig. S4**), and were similar between *SORL1* WT-AD (n=78; mean: 462 ± 134 pg/ml) and *SORL1* WT-NC (n=50; mean: 474 ± 131 pg/ml) (p=0.45) (**Table S2**), such that these groups were merged into a single “*SORL1 WT*” control group for further analysis. Additionally, due to limited statistical power, carriers of 2 or 3 NPVs or 2 MPVs were grouped according to the highest predicted effect for further analysis.

We previously established that SORL1 variants adhering to different priority groups associated with specific effects on AD risk (i.e. odds ratios): OR_PTV_ = 17.2, OR_HPV_ = 6.1, OR_MPV_ = 1.5, OR_LPV_ = 1.2, OR_NPV_ = 1.1. This then allowed us to investigate variant-associated sSORLA concentrations relative to category-specific AD-risk. We found that carriers of variants with higher predicted risk for AD, have lower sSORL1 concentrations: for each 1-unit increase in the natural log-transformed odds ratio (ln(OR)), sSORL1 concentrations decreased by 87.4 pg/ml (β = −87.4; SE = 12.4; p=1.9×10^−12^)(**Fig S5**).

Next, we compared CSF-sSORL1 concentrations between individual variant groups and *SORL1* WT controls (**Fig 3** and **Table 2**). The average CSF-sSORL1 concentration across the 128 WT-*SORL1* carriers was 466 pg/ml (SD = 133). As expected PTV carriers, who carry only one functional allele had significantly lower CSF-sSORL1 concentrations compared to the group of SORL1 WT carriers (i.e the merged SORL1 WT AD cases and controls) (260 pg/ml ± 123 SD, p=2.1×10^−7^). Notably, CSF-sSORL1 concentrations of high-priority missense variants were lowest of all prioritized missense variant groups, and significantly lower than the SORL1 WT controls (323 pg/ml ± 154 SD, p=9.8×10^−8^). Specifically, recent functional studies on the Y1816C and D1105H variants have demonstrated their pathogenicity and causative effect on AD (17, 36): in CSF of the carriers of these variants, we observed very low sSORL1 concentrations (305 pg/ml for Y1816C; 270 and 309 pg/ml for D1105H). In contrast, the sSORL1 concentrations of MPV, LPV, and NPV carriers were not significantly different from the WT controls (p_adj_ >0.00625). In-frame indels, for which AD risk estimates are currently unavailable and which cannot be assigned to any prioritization group, showed significantly reduced concentrations compared to WT controls (315 ± 75 pg/ml, p=1.1×10^−3^). Finally, CSF-sSORL1 concentrations in carriers of the E270K and A528T common variants were not significantly different compared to WT controls (E270K: 420 pg/ml ± 135 SD, p = 0.29; A528T: 464 pg/ml ± 118, p=0.72).

**Table 2.**
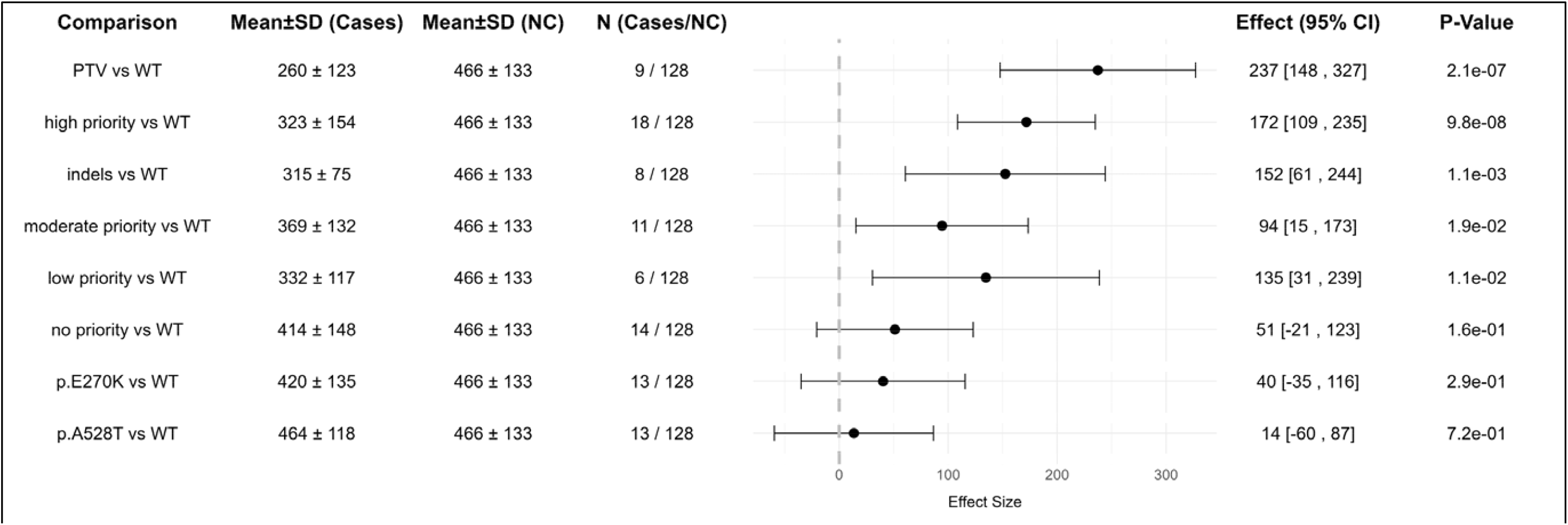
The effect of having a specific SORL1 variant on the CSF-sSORL1 concentration, as determined by a robust linear regression for CSF-sSORL1 concentrations between the different severity groups versus Wild-type (*SORL1* WT AD cases + controls combined). WT = Wild-type, SD = Standard Deviation, CI = Confidence Interval.

**Fig 3.**
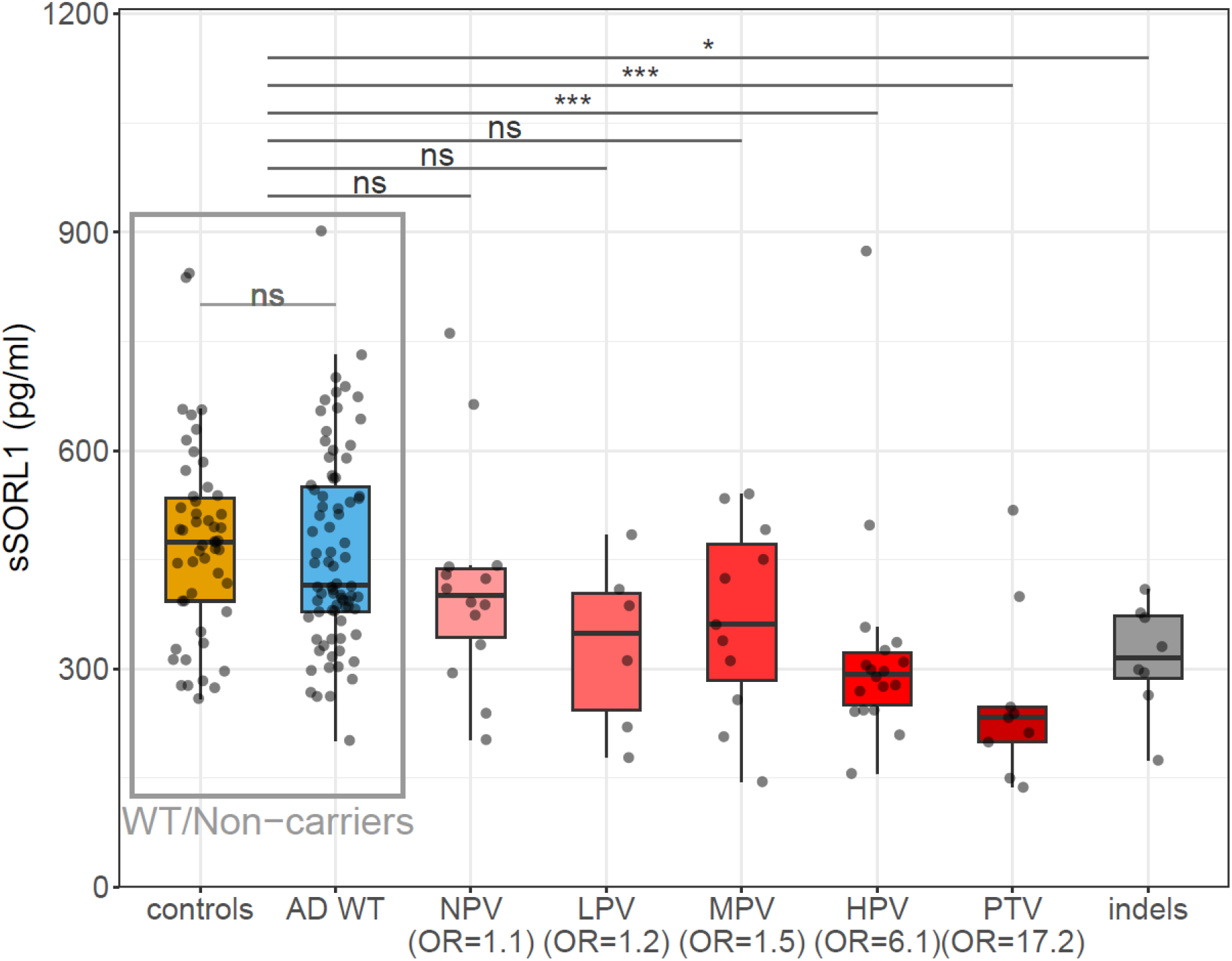
sSORL1 concentrations vs pathogenicity category. ELISA measurements of sSORL1 concentrations in CSF derived from carriers of *SORL1* genetic variants with different pathogenicity categories or carriers of WT *SORL1*. Sample size per group: Controls = 50, AD WT= 78, NPV = 14, LPV = 6, MPV = 11, HPV = 18 and PTV = 9. Abbreviations: HPV = High priority variants; indels = in-frame insertion deletion variant; LPV = Low priority variant; MPV = moderate priority variant; NPV = No priority variant; OR = Odds ratio; PTV = Protein truncating variant; WT = Wild type. Adjusted significance threshold: 0.000625 <p≤0.00625 = *; 0.0000625 <p≤0.000625 = **; p≤0.0000625 = ***.

The distribution of sSORLA concentrations was wide, prompting us to estimate the lower bound (lowest 1%) of the CSF-sSORL1 concentration range in WT *SORL1* controls. We found the lower-bound CSF concentrations to be at 260 pg/ml (95% CI: 202–274 pg/ml) sSORL1 concentrations below this lower-bound extreme can be considered abnormally low. Notably, 12 out of 27 (44%) HPV and PTV carriers had sSORL1 levels below this threshold. In addition, several carriers of variants within the non-pathogenic WT, MPV, LPV and, NPV categories had very low sSORL1 levels. Specifically, while variants translating to G1440R and S1107N, affecting the CR domain, were conservatively classified as a ‘moderate priority variant’ due to a lack of evidence for pathogenicity by DMDM analysis (19), our results suggest that these variants may have an impairing effect on SORL1 function. On the other hand, we observed that several HPV and PTV carriers had unexpectedly high sSORL1 levels. For example, we observed an outlier high sSORL1 concentration in the CSF derived from a carrier of an exon 13-splicing variant that likely causes an exon skipping frameshift, expected to lead to nonsense mediated RNA decay. Possibly, the incorrect splicing may be incomplete such that the haploinsufficiency observed for PTVs is less pronounced. Alternatively, in addition to impairing genetic variation, sSORL1 levels may be driven by alternative, currently unknown mechanisms.

### Measurement of sSORL1 in CSF and cell culture media using Western-Blot and sSORL1 in plasma

We validated ELISA-based sSORL1 measurements in CSF using WB in a subset of 37 individuals, observing a strong correlation between the two methods (ρ = 0.541; p = 6.6 × 10^−6^; **Fig S6**). WB images and comparisons across pathogenicity categories are provided in the **Supplement (Fig. S7 and S8;** uncropped blots of S7 are included as Supplementary File).

In agreement with ELISA and WB, a cellular model expressing either high-priority missense variants or in-frame indels exhibited significantly reduced sSORL1 secretion compared to WT, also these models show no difference in sSORL1 secretion in media between cells expressing the common E270K and A528T variants and those expressing WT SORL1 (**Supplemental Methods, Fig. S9**; uncropped blots of S9 are included as Supplementary File). Finaly, for 44 individuals, both CSF and plasma samples were available, which indicated that plasma sSORL1 concentrations did not correlate with CSF sSORL1 concentrations (ρ = 0.126; **Fig. S10**). A comparison between pathogenicity categories, is provided in the Supplement (**Fig. S11**).

### Association of CSF sSORL1 concentrations with APOE genotype, MMSE, age of diagnosis, p-Tau181, t-Tau and A-beta 42

Lastly, we investigated whether CSF-SORL1 levels in *SORL1* WT AD patients were significantly associated with clinical measures of dementia and other CSF-biomarker levels, including *APOE*-ε4 genotype, MMSE, age, sex, Aβ42, tTau, and pTau-181, accepting associations with p_adj_ <0.00714 as significant. CSF-sSORL1 levels were not significantly associated with age at AD diagnosis (β = 3.4; SE = 2.7; *p* = 0.21) (**Fig. 4A**). Furthermore, we observed that females trended to have lower CSF-sSORL1 levels (mean = 437 pg/ml) compared to males (mean = 511 pg/ml), although with a *p*-value of 0.033 this was not accepted as a significant correlation (**Fig. 4B**). Similarly, higher CSF-sSORL1 levels trended to be associated with lower cognitive performance as measured by MMSE score, but this did not reach significance (β = –6.3; SE = 3.4; *p* = 0.06) (**Fig. 4C**). CSF-sSORL1 did not associate with *APOE ε4* genotype (β = –7.6; SE = 20.6; *p* = 0.71; data not shown), and we observed no significant interaction effect between *APOE ε4* status and *SORL1* pathogenicity category (p = 0.19). While CSF-sSORL1 did not associate with CSF-Aβ42 concentrations (β = 0.03; SE = 0.06; *p* = 0.62) (**Fig. 4D**) we observed a positive correlation with both tTau (β = 0.15; SE = 0.04; *p* = 5.8×10^−4^) and pTau-181 (β = 1.8; SE = 0.41; *p* = 9.7×10^−6^), which seems to be driven by the subset of AD patients with the highest levels of CSF-(p) and (t)Tau (**Fig. 4E** and **F**). Then, we investigated to what extent correlations between CSF-sSORL1 concentrations and AD biomarkers covered the wider cognitive spectrum. Upon correlating CSF-sSORL1 across the union of AD cases and controls who were sSORL1-WT, we observed only a weak trend for an (unexpected) positive association with CSF-Aβ42 concentrations (p=0.022), and while the correlation with pTau-181 was weaker than when conditioning on AD patients only, it remained significant (p=3.1×10^−3^) (**Fig. S12**).

**Fig 4.**
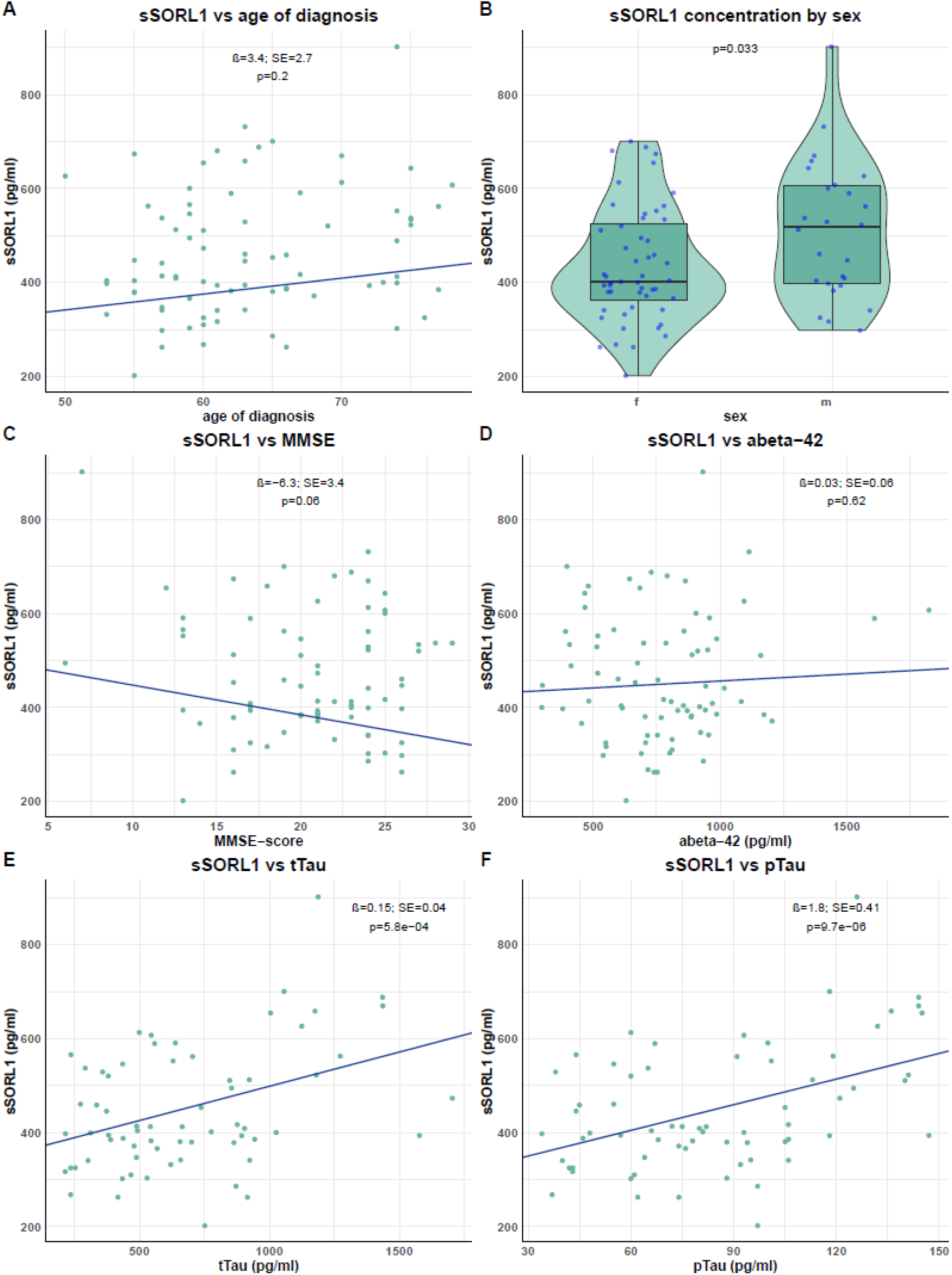
Correlation of sSORL1 levels SORL1 WT AD patients (n=78), with age of diagnosis, sex, MMSE and AD biomarkers. To assess the relationship between CSF-sSORL1 concentrations and age at diagnosis (A), sex (B). MMSE (C) and AD biomarkers (D-F) in SORL1-WT AD, we performed a Robust Linear Model to account for the potential influence of outliers. The association of age of diagnosis with CSF-sSORL1 was corrected for sex and APOE-status. The association of sex with CSF-sSORL1 was corrected for age of diagnosis. The association of MMSE and the AD biomarkers with CSF-sSORL1 were corrected for age of diagnosis and sex.

## Discussion

CSF-sSORL1 concentrations were strongly variable, and distributions overlapped between AD patients and healthy controls. However, CSF-sSORL1 concentrations in carriers of high-priority *SORL1* missense variants (HPVs) were significantly lower than those observed in *SORL1* WT AD patients or controls, and resembled concentrations observed in carriers of protein-truncating variants (PTVs). With this, we identify CSF-sSORL1 concentrations as a promising candidate biomarker to support variant-specific functional pathogenicity in carriers of a possibly pathogenic *SORL1* variant.

sSORL1 concentrations in CSF derived from carriers of a *SORL1* HPV represent the extreme lower end of the sSORL1-concentration distribution. Indeed, we observed very low CSF-sSORL1 concentrations in carriers of the mechanistically proven pathogenic Y1816C and D1105H variants (17, 36, 37). This substantiates the findings by Castelot et al. who used WB to show that CSF-sSORl1 concentrations in carriers of ‘trafficking defective’ variants are significantly decreased (21). Furthermore, we observed reduced levels of sSORL1 in carriers of PTVs, which is fully consistent with expectations for losing one copy. Further, sSORL1 concentrations in carriers of in-frame indel variants were significantly reduced compared to those observed in carriers of WT-*SORL1* but had slightly higher CSF-sSORL1 concentrations than PTV- and HPV-carriers. While this possibly represents a milder, yet impairing, effect of the tested in-frame variants on SORL1 function compared to PTVs/HPVs, further functional investigation is necessary. Moreover, carriers of several MPVs had very low CSF-sSORL1 concentrations, suggesting that despite the lack of predicted pathogenicity using DMDM, these variants may have an impairing effect on SORL1 function. On the other hand, for some HPV and PTV we observe relatively high CSF-sSORLA levels. Together, it is likely that factors other than rare coding genetic variants influence sSORL1 concentrations in CSF, this may concern non-coding genetic and epigenetic variations associated with different *SORL1* haplotypes (38), but also, given their trends for association, effects of sex and cognitive performance (i.e. disease severity).

Indeed, when considering CSF-sSORL1 levels in individuals who are WT for SORL1, previous studies have reported conflicting results regarding associations with AD. Using a WB approach, Ma et al. observed *decreased* CSF-sSORL1 concentrations in 13 AD patients with early changes of AD-associated cognitive decline (mean MMSE: 24; mean age at onset 74y) relative to 13 controls (39). Ikeuchi et al. developed an ELISA assay and found *increased* CSF-sSORL1 levels in 29 AD patients with more severe cognitive decline (mean MMSE 15.4; mean age at onset 64y) relative to 27 controls (40). A more recent study using WB by Castelot et al. found no significant difference in CSF-sSORL1 concentrations between 40 AD patients (mean age at onset: 60y) and 30 controls (21). In our sample, CSF-sSORL1 concentrations did not distinguish between 78 amyloid-positive SORL1 WT AD cases with intermediate AD severity (median MMSE 20.7; median age at onset 63y) and 50 controls. Our results are therefore in line with those reported by Castelot et al., suggesting that CSF-sSORL1 levels may not consistently differ between AD patients and controls. While differences between used CSF-sSORL1 assays may contribute to these discrepancies, future research will have to test to what extent CSF-sSORL1 concentrations are affected by age, sex or with disease severity.

Upon further investigation of associations between CSF-sSORL1 levels and aspects of AD, we found that CSF-sSORL1 concentrations did not correlate with CSF-Aβ42 concentrations *within* our sample of amyloid positive *SORL1* WT AD patients, despite SORL1’s role in APP processing and Aβ degradation (6-11). Thus, while CSF-Aβ42 clearly distinguishes between AD cases and controls (25), CSF-Aβ42 concentrations may not contribute additional meaningful signal within advanced (amyloid positive) AD cases, which might preclude any association with CSF-SORL1. In contrast, we observed a positive association between CSF-sSORL1 with (p)Tau concentrations within the SORL1-WT AD cases. Since elevated CSF (p)Tau levels reflect an increased abundance of neuropathological amyloid plaques and neurofibrillary tangles (25, 41), we speculate that elevation of CSF-sSORL1 concentrations may reflect overall increased retromer activity in which the cell attempts to rescue the cell stress brought about by AD-associated processes. In addition, a recent report provided evidence that SORL1 serves as a binding partner for tau, such that as a consequence of SORL1’s elevated expression, it may promote tau seeding, thereby contributing to the spreading of tau pathology and further aggravating the disease process (42). Notably, this observation might explain why the association between CSF-sSORL1 and CSF-(p) tau levels seems to be driven by the subset of AD patients with the highest CSF-(p) tau levels, which is not captured by AD/control comparison. While the cross-sectional design of our study precluded us from assessing fluctuations in CSF-sSORL1 concentrations with different disease stages, this should clearly be an important objective for future studies.

Here, we provide the first thorough technical validation of an ELISA assay for sSORL1 in CSF. The ELISA detects the SORL1 VPS10p domain. No nonspecific signals were detected in *SORL1* knockout (KO) cells, and the assay showed no cross-reactivity with other VPS10p family members (SORT1 and SORCS2). However, since the epitopes of the antibodies are not fully characterized, we cannot exclude that binding may depend on a properly folded VPS10p domain. While calibrated with a recombinant SORL1 VPS10p fragment, the assay successfully detected full-length sSORL1. Importantly, CSF-sSORL1 concentrations were consistent between ELISA and WB analyses across genetic variant carriers, reinforcing the robustness of sSORL1 detection in CSF.

The investigation of CSF-SORL1 concentrations in multiple individuals who carry the same rare mutation contributes to an increased understanding of robustness of specific variant-effects on in CSF-sSORL1 (17, 36), but the extreme rarity of impairing SORL1 variants precludes such an analyses for most. Here, we presented five carriers of the p.F1123-R1124delinsLS variant (four AD patients and one SCD patient), in whom we observed consistent CSF-sSORL1 concentrations, suggesting a robust variant-specific effect on CSF-sSORL1 levels. Furthermore, we found that variants, grouped according to having a similar predicted effect on AD risk, have a similar effect on CSF-sSORL1 concentrations, further supporting the observation that genetic SORL1 impairment may have a robust variant-specific effect on CSF-sSORL1 concentrations.

In conclusion, our findings suggest that CSF-sSORL1, as quantified by the sSORL1 ELISA assay, may represent a valuable biomarker for confirming the pathogenicity of specific rare *SORL1* variants. Despite strong inter-individual variability in CSF-sSORL1 concentrations, our results show a robust decrease in CSF-sSORL1 concentrations in carriers of rare impairing *SORL1* missense variants, comparable to the decrease observed in carriers of PTVs. Further refinement of the clinical utility of CSF-sSORL1 in confirming variant pathogenicity is warranted, including further investigations on the impact of factors including disease status and sex on the wide variance in concentrations. Together, we hypothesize that genetic impairment of SORL1 leads to increased risk of AD due to the inability of the retromer to traffic cargo and to rescue early AD-associated cell stress, predisposing variant carriers to the development of advanced AD processes. Whether on the other hand, enhanced SORL1 expression may contribute to increased tau seeding and advancement of AD-associated processes will have to be focus of future studies in larger cohorts with longitudinal design.

## Declarations

### Ethics approval and consent to participate

All patients gave informed consent for the use of their medical data and biomaterial. The study was approved by the Medical Ethics Committee of Amsterdam UMC and conducted in accordance with the Declaration of Helsinki.

### Consent for publication

Not applicable

### Availability of data and materials

The data supporting the findings of this study are not publicly available due to sensitivity concerns. However, they can be accessed upon reasonable request from the corresponding author, subject to approval by the Medical Ethics Committee of Amsterdam UMC.

### Author contributions

Conceived the study: OA, CET, HH;

Wrote the manuscript: MdW, SvdL, JR, LV, MH, DB, WF, OA, CET, HH;

Data acquisition: MdW, SvdL, ML, LB, NB, GM, AMJ, CV, SC, JS, CT, MMW, RvS, GO, DJ, AM, DB;

Analyses: Mdw, SvdL, ML, LB, NB, GM, LV, NT, MH, WF, OA, CET, HH;

All authors read, revised and approved the final manuscript.

## Acknowledgments

We thank all study participants and all personnel involved in data collection for the contributing studies. Research of Alzheimer center Amsterdam is part of the neurodegeneration research program of Amsterdam Neuroscience. Alzheimer Center Amsterdam is supported by Stichting Alzheimer Nederland and Stichting Steun Alzheimercentrum Amsterdam. We also acknowledge Søren Thirup (Aarhus University) for providing the sortilin plasmid.

