## Supplemental data for "Soluble SORL1 in cerebrospinal fluid as a marker for functional impact of rare SORL1 variants"

TO:

**Index**

Supplemental methods

### WES analysis

Exome sequencing for the analysis was previously described (1). In short, the exomes from the ADC-VUmc cohort were captured with the Nimblegen SeqCap EZ Exome capture kit v2/v3/MedExome, KAPA Hyper, Twist Comprehensive or the Agilent V6. DNA from all samples was prepared with the Illumina TruSeq Paired-End Library Preparation Kit and 100 bp paired-end reads were acquired by sequencing the libraries on a HiSeq 2000 or 2500. We sequenced to at least 40x mean coverage to ensure sufficient read depth for variant calling. For *SORL1* variant calling, the intersection of the capture kits covered 93.2% of exons 1-47 and 2.7% of exon 48 in the *SORL1* gene. Genotype calling was done using holstege 2022 et al. pipeline, as decribed previously. In addition, new samples were processed using a standardized pipeline, employing DeepVariant 1.6.1 (2) as the variant caller and dragmap-1.4 as aligner. The threshold for variant calling was >20 for genotype quality and >10 for the total reading depth per variant (as outputted by the variant caller). The missing rate for the called variants was 0.51% on average.

### Analytical validation

We performed analytical validation following international protocol as described by Andreasson et al. (3).

#### Parallelism

For parallelism, samples (n=5) were diluted using 2-times serial dilution. Dilutional linearity was tested by spiking samples (n=3) with recombinant SORL1 (3000 pg/ml), which were serially diluted (dilution factor of 3). The recovery % was calculated by ((concentration at dilutionfactor X*dilution factor X)/(concentration of previous dilution factor*previous dilution factor)*100). Recovery was assessed by diluting samples 10-times and afterwards spiked with several concentrations of recombinant SORL1 (2500, 300, 65, 0 pg/ml). To calculate the % recovery, the following formula was used: (concentration spiked samples – concentration neat sample) / theoretical concentration spiked samples * 100.

#### Sample stability

Sample stability was evaluated by observing the effect of freeze/thawing on the sSORL1 concentrations in samples (n=3) which were exposed to a series of 4 freeze/thaw cycles. The sSORL1 concentration of the samples was normalized to the reference condition.

#### Specificity/cross-reactivity

Specificity/cross-reactivity was assessed by performing three different experiments. Firstly, to test specificity, we determined to what extent the ABCAM ELISA picked up increased concentrations of other proteins from the VPS10-family; namely, the SORT1 protein (see Methods: Recombinant SORT1 protein) and commercially available SORCS2 protein (R&D systems, 4238-SR). We tested four concentrations of recombinant SORL1 (0 pg/ml, 125 pg/ml, 500 pg/ml, and 2000 pg/ml), and samples were spiked with varying concentrations of recombinant SORT1 or SORCS2 (0 pg/ml, 125 pg/ml, 500 pg/ml, 2000 pg/ml). Second, we tested full size recombinant SORL1 to see if the assay is able to detect the full protein and not just a fragment. For this experiment, we generated recombinant SORL1 (see Methods: recombinant sSORL1) and a commercially available recombinant SORL1 protein (R&D Systems, 11083-LA). Blank samples were spiked with four different concentrations of the in-house generated or commercial full-length recombinant SORL1 proteins (0 pg/ml, 125 pg/ml, 500 pg/ml, and 2000 pg/ml). Additionally, to test for possible interference a sample containing 500 pg/ml of the standard recombinant SORL1 protein (recombinant from the ABCAM ELISA) was spiked with either 500 or 2000 pg/ml of both full-length recombinant SORL1. Finally, to determine whether non-specific signal was present, we measured the sSORL1 concentrations in SORL1 knock out and SORL1 WT iPSCs (see below for Methods). For this, we used mock cells as a positive control, which underwent the entire CRISPR processing workflow, but without any genetic modification (see below). To investigate potential non-specific signal across different cellular contexts, we included 20-day-old induced neurons, 15-day-old cortical organoids, and 60-day-old cortical organoids. Protein levels were stable over time in neurons, so only one neuronal time point was used, while organoids showed variation over time, warranting inclusion of both early and late stages.

To assess potential non-specific signal across different cellular contexts, SORL1 KO samples were analyzed from 20-day-old induced neurons, known for high SORL1 expression in wild-type conditions, as well as from 15-day-old and 60-day-old cortical organoids to capture early and mature stages of organoid development

#### Estimation of sSORL1 concentration in CSF

Each pair of CSF samples from a *SORL1* variant carrier and the matched wild-type AD patient was run on a separate gel. To prepare the standard curve, different dilutions of sSORL1 recombinant protein ranging from 125 to 12.5 ng/mL were used (see methods describing the generation of the sSORL1 recombinant protein. CSF samples from the *SORL1*-variant carrier and the matched *SORL1*-WT AD patients were diluted 3:4 and 1:2 in loading buffer, and each diluted condition was run in duplicate. The pair of CSF samples was loaded blinded, and the experimenter was not informed about the pathogenicity group to which the *SORL1*-variant carried by the CSF sample donor belonged. sSORL1 concentration in the CSF samples was estimated by annotating the mean of the band signals corresponding to the 3:4 and 1:2 dilutions of each sample. For samples for which the signal for one of the two dilutions fell outside the linear range of the standard curve, we estimated the sSORL1 concentration for the samples using only the one dilution within the linear range. The standard sSORL1 signal varied across individual gels, which led to too much uncertainty for the estimation of *absolute* sSORL1 concentrations. Instead, we estimated the *rel-sSORL1* concentrations by comparing the signal for each unique *SORL1* variant carrier with that of the matched sample (set to 100%) from individual gels.

### CRISPR/Cas9 knock-out of SORL1 in iPSCs and organoids

#### i3N induced pluripotent stem cell line cultivation

Two cell lines of human induced pluripotent stem cells (iPSCs) were used in this study. Specifically, i3N iPSCs with stably integrated doxycycline-inducible NGN2 (RRID:CVCL_C7XJ), generously provided by Michael Ward (4) and CTRL-E3 iPSCs (UIOi002-A (RRID:CVCL_C1X2)), generously provided by Li-Huei Tsai (5). Doxycycline-inducible NGN2 transgene (Addgene # 127288) was stably integrated into this cell line upon lentiviral transduction (Raska et al. in preparation). Both cell lines were propagated as a feeder-free monolayer culture on Matrigel® hESCs-qualified matrix (Corning) in mTeSR™1 medium (STEMCELL Technologies) and passaged using TrypLE™ reagent (Gibco).

#### Generation of CRISPR/Cas9 knock-out of SORL1

The generation of iPSCs with a SORL1 knock-out (KO) was achieved using CRISPR/Cas9, as described in Jensen et al., 2024 (6). In brief, the SORL1-KO was introduced into i3N iPSCs and CTRL-E3 iPSCs through an RNP  (ribonucleoprotein) complex consisting of Cas9-GFP (Sigma) and a previously published gRNA targeting exon 6 of the SORL1 gene (7). The RNP complexes were delivered into cells via the Neon™ Transfection System (Thermo Fisher Scientific), followed by sorting transfected cells based on the GFP signal to enhance CRISPR efficiency. For SORL1 KO generation, cells were single-cell sorted into 96-well plates, and the resulting clones were screened for the absence of SORL1 protein expression using Western blotting (LR11 antibody, BD Tranduction Lab; 611861). Final verification of SORL1 KO clones was performed via Sanger sequencing (Seqme). To ensure that observed effects were not attributable to procedural modifications, we generated Mock cells that underwent the entire CRISPR processing workflow without any actual genetic modification. This included cell sorting and cultivation as single-cell clones to replicate the selective bottleneck imposed by the procedure.

#### 2D neuronal cultivation and sample collection

For the differentiation of i3N iPSCs into neurons, we used a previously published protocol (4). Briefly, undifferentiated iPSCs from both cell lines (Day 0) were plated as a single-cell monolayer on a Matrigel® coated dish at the density of 0.7x105 cells/cm2 into an Induction medium (IM) supplemented with 2 μg/mL doxycycline (Sigma) and 10 μM Y-27632 ROCK inhibitor (Selleckchem). For 2 days (D2-3), IM with 2 μg/mL doxycycline was changed daily. At D3, cells were re-plated onto a Poly-L-Ornithine (Sigma) coated dish with Cortical Neuron medium (CNM) at the density of 0.8x105 cells/cm2. Half of the CNM medium was changed twice a week. Before sample collection, CNM media was fully changed for Essential 6 medium (Gibco) and cells were cultivated for 3 days. All cell media samples and the whole cell lysate samples were collected on day 20. Cell media samples were supplemented with cOmplete™ Mini protease inhibitor (Roche), stored at -80°C and subsequently used for ELISA. Whole cell lysate samples were lysed on ice for 1 hour in lysis buffer (1M Tris HCl pH 8.1, 0.5M EDTA, 1% Triton-X-100, 1% NP-40) containing cOmplete™ Mini protease inhibitor (Roche). Total protein concentration in lysates was then determined using the DC-assay kit (BioRad) according to the manufacturer’s instructions and used to normalize ELISA.

#### 3D cerebral organoid cultivation and sample collection

3D cerebral organoids were differentiated as previously described in Vanova et al., 2023 (8). Briefly, both iPSC lines were detached by Accutase (Thermo Fisher Scientific) and plated at day 0 (D0) into non-adherent V-shaped 96-well plates at the density of 2,000 cells per well in mTeSR™1 medium (STEMCELL Technologies) with 50 μM ROCK inhibitor (Y-27632, Selleckchem) to induce embryoid body (EB) formation. When EBs reached the size of at least 400 μm, a fresh Neural-Induction Medium was added to EBs every day for 6 days (usually from D3 to D8). Following this period, organoids were embedded in 7 μL of cold Geltrex (Thermo Fisher Scientific). Solidified Geltrex droplets with organoids were gently detached and cultured without shaking in the Cerebral Organoid Differentiation Medium (CODM) without vitamin A for 7 days. Subsequently, organoids were cultured in CODM with vitamin A and moved to an orbital shaker at D24-D28. Cell media samples and whole cell lysate samples were collected on day 15 and 60. Cell media samples were supplemented with cOmplete™ Mini protease inhibitor (Roche) and stored at -80°C. Whole-cell samples were lysed on ice for 1 hour in lysis buffer (1M Tris HCl pH 8.1, 0.5M EDTA, 1% Triton-X-100, 1% NP-40) containing cOmplete™ Mini protease inhibitor (Roche) and used for ELISA. Total protein concentration in lysates was determined using the DC-assay kit (BioRad) according to the manufacturer’s instructions and used to normalize ELISA.

### Recombinant proteins

#### Recombinant sSORL1 protein

The part of SORL1 corresponding to the fragment secreted to the CSF (residues 54-2107) was prepared as previously described by Jacobsen et al. 2001 (9). In brief, CHO-K1 cells were cultivated in HyQ-CCM5 medium (HyClone, Logan, UT) devoid of serum, and transfected with a pcDNA 3.1/Zeo(+) vector containing the cDNA encoding for the luminal part of SORL1 using FuGENE 6 Transfection reagent according to manufacturer’s instructions (Promega, cat# E2691). Selection of stably transfected cells was carried out with Zeocin-containing medium at a concentration of 500 mg/mL (InvivoGen, #ant-zn-05). sSORL1 was purified from the medium of transfected cells using RAP affinity column as described. In addition to the generated recombinant sSORL1 protein, we also used a commercially available recombinant SORL1 protein (R&D Systems, 11083-LA).

#### Recombinant SORT1 protein

Human sortilin, encoding residues 1 to 723, followed by a C-terminal hexa-histidine tag, was expressed in FreeStyle HEK 293F cells (Invitrogen) and purified using IMAC, as previously described in Januliene et al, 2017 (10). Briefly, the harvested conditioned media was buffered with 50 mM Tris-HCl pH8.0 and applied to Ni-NTA column, equilibrated in 50 mM Tris HCl pH 8.0, 150 mM NaCl. The column was washed with five column volumes of the same buffer, additionally containing 10 mM imidazole and eluted with the buffer, containing 250 mM imidazole. Sortilin was further purified via SEC, using Superdex 200 10/300 GL column (GE Healthcare), concentrated to 1.4 mg/ml and stored at -80°C until use.

#### Recombinant SORCS2 protein

Experiments were performed using commercially available recombinant human SORCS2 protein (R&D Systems, 4238-SR).

### Neuroblastoma cell lines

#### Neuroblastoma cell lines

Mouse neuroblastoma N2a cells were cultivated in Dulbecco’s modified Eagle’s medium (DMEM, Sigma) supplemented with 10% fetal bovine serum and penicillin/streptomycin. 5 x 10^5^ cells were transfected in duplicate with constructs encoding for SORLA-WT, SORLA-D1105H, SORLA-Y1816C, SORLA-FR1123/1124LS, SORLA-E270K, or SORLA-A528T using FuGENE HD Transfection Reagent according to manufacturer’s instructions (Promega, #E2311) (see supplement for construct-containing plasmids). 48 hs after transfection, cell medium was changed to serum-free conditional medium, and following additional 48 hs both cell lysates and media were harvested. Four independent experiments of cell transfection were performed for each construct.

Equal amounts of lysates and media samples were separated by SDS-PAGE and subsequently transferred to nitrocellulose membranes. After blocking (as described above in WB section), membranes were incubated with primary antibody overnight at 4°C (mouse anti-LR11; mouse anti-actin, Sigma, #A2066). Quantification of sSORLA signal was carried out using ImageJ software and the results were plotted on a graph as mean of duplicate values relative to the mean signal for SORLA-WT samples for each of four independent experiments (N=4).

#### Primers:

The expression construct for the full-length SORLA-WT receptor was previously described (9), and used as template for the introduction of the *high priority* variants D1105H and Y1816C, the *in-frame indel* FR1123-1124LS variant, and the common variants E270K and A528T by site-directed mutagenesis following manufacturer’s instructions (Agilent). The following primers were used to insert the desired mutations: D1105H fw 5’-CTTTGACAACGACTGTGGACACATGAGCGATGAGAGAAAC-3’, and D1105H rev 5’-GTTTCTCTCATCGCTCATGTGTCCACAGTCGTTGTCAAAG-3’; FR1123-1124LS fw 5’-GACCTGGACACCCAGTTAAGTTGCCAGGAGTCTGGG-3’, and FR1123-1124LS rev 5’-CCCAGACTCCTGGCAACTTAACTGGGTGTCCAGGTC-3’; Y1816C fw 5’-GGCAATCTGACAGCTCATACATCCTGTGAGATTTCTGCCTGGGCCAAGACTG-3’, Y1816C rev 5’-CAGTCTTGGCCCAGGCAGAAATCTCACAGGATGTATGAGCTGTCAGATTGCC-3’; E270K fw 5’-CATCTACATTGAACGACATAAACCCTCTGGCTACTCCACTG-3’, E270K rev 5’-CAGTGGAGTAGCCAGAGGGTTTATGTCGTTCAATGTAGATG-3’; A528T fw 5’-CTCTAGCAGTGCTGGAACCAGGTCGAGAGGCAC-3’, A528T rev 5’-GTGCCTCTCGCCACCTGGTTCCAGCACTGCTAGAG-3’.

All mutated constructs were verified by Sanger sequencing (Eurofins Genomics).

Supplemental tables

### Supplemental table 1. Variants in CSF-cohort by pathogenicity category.

| Pathogenicity category | variant | domain | p. cons | mutation | CADD | revel score | | EOAD OR | LOAD OR | WB rel levels | elisa conc. |
| --- | --- | --- | --- | --- | --- | --- | --- | --- | --- | --- | --- |
| PTV | chr11:chr11:121520673:  CCATTT>AAATGCAAATGAAGTCAGCAAA | N/A | p.P410Kfs*4 | stopgain | 35 | | N/A | 35.31 | 8.57 | 74 | 518.124 |
| PTV | chr11:chr11:121543728:T>C | N/A | NA | splicing | 25.7 | | N/A | 35.31 | 8.57 | 108 | 399.503 |
| PTV | chr11:chr11:121557338:C>T | N/A | p.R866X | stopgain | 41 | | N/A | 35.31 | 8.57 | 21 | 233.177 |
| PTV | chr11:chr11:121559608:G>GCTCA | N/A | p.G1003Hfs*37 | frameshift | 31 | | N/A | 35.31 | 8.57 | 52 | 247.704 |
| PTV | chr11:chr11:121604292:CCTG>GCTCGGAC | N/A | p.E1543Gfs*5 | frameshift | 0 | | N/A | 35.31 | 8.57 | 73 | 238.846 |
| PTV | chr11:121558808:CCGCA>C | N/A | p.H962Pfs*45 | frameshift | N/A | | N/A | 35.31 | 8.57 | N/A | 137.484 |
| PTV | chr11:121558808:CCGCA>C | N/A | p.H962Pfs*45 | frameshift | N/A | | N/A | 35.31 | 8.57 | N/A | 150.057 |
| PTV | chr11:121595670:C>T | N/A | p.R1473X | stopgain | N/A | | N/A | 35.31 | 8.57 | N/A | 199.673 |
| PTV | chr11:121595691:C>T | N/A | p.Q1480X | stopgain | 37 | | N/A | 35.31 | 8.57 | N/A | 212.573 |
| HPV | chr11:chr11:121543604:T>G | VPS10p | p.V581G | nonsynonymous SNV | 28 | | 0.542 | 9.93 | 4.2 | 178 | 325.731 |
| HPV | chr11:chr11:121543625:C>T | VPS10p | p.T588I | nonsynonymous SNV | 32 | | 0.562 | 9.93 | 4.2 | 45 | 241.583 |
| HPV | chr11:chr11:121543706:T>G | VPS10p | p.V615G | nonsynonymous SNV | 28.9 | | 0.629 | 9.93 | 4.2 | 186 | 336.592 |
| HPV | chr11:chr11:121550590:G>A | CC | p.R729Q | nonsynonymous SNV | 31 | | 0.792 | 9.93 | 4.2 | 67 | 243.464 |
| HPV | chr11:chr11:121570246:G>C | CR | p.D1105H | nonsynonymous SNV | 31 | | 0.96 | 9.93 | 4.2 | 35 | 269.567 |
| HPV | chr11:chr11:121570246:G>C | CR | p.D1105H | nonsynonymous SNV | 31 | | 0.96 | 9.93 | 4.2 | 53 | 309.642 |
| HPV | chr11:chr11:121614898:A>G | 3FN | p.Y1816C | nonsynonymous SNV | 31 | | 0.897 | 9.93 | 4.2 | 30 | 305.356 |
| HPV | chr11:121496993:G>A | VPS10p | p.E295K | nonsynonymous SNV | 27.9 | | 0.53 | 9.93 | 4.2 | N/A | 874.192 |
| HPV | chr11:121514282:A>G | VPS10p | p.Y391C | nonsynonymous SNV | 27.3 | | 0.477 | 9.93 | 4.2 | N/A | 497.631 |
| HPV | chr11:121550082:C>T | CC | p.T725M | nonsynonymous SNV | 25.4 | | 0.653 | 9.93 | 4.2 | N/A | 357.386 |
| HPV | chr11:121550604:G>A | CC | p.D734N | nonsynonymous SNV | 31 | | 0.6 | 9.93 | 4.2 | N/A | 243.524 |
| HPV | chr11:121550604:G>A | CC | p.D734N | nonsynonymous SNV | 31 | | 0.6 | 9.93 | 4.2 | N/A | 296.989 |
| HPV | chr11:121554078:A>G | YWTD | p.Y803C | nonsynonymous SNV | 28.9 | | 0.934 | 9.93 | 4.2 | N/A | 156.423 |
| HPV | chr11:121570171:C>T | CR | p.R1080C | nonsynonymous SNV | 28.3 | | 0.602 | 9.93 | 4.2 | N/A | 289.258 |
| HPV | chr11:121570241:G>A | CR | p.C1103Y | nonsynonymous SNV | 29.1 | | 0.963 | 9.93 | 4.2 | N/A | 209.561 |
| HPV | chr11:121574273:C>T | CR | p.R1124C | nonsynonymous SNV | N/A | | 0.858 | 9.93 | 4.2 | N/A | 299.143 |
| HPV | chr11:121574273:C>T | CR | p.R1124C | nonsynonymous SNV | N/A | | 0.858 | 9.93 | 4.2 | N/A | 275.995 |
| HPV | chr11:121591058:A>G | CR | p.Y1424C | nonsynonymous SNV | 22.3 | | 0.6 | 9.93 | 4.2 | N/A | 277.93 |
| indels | chr11:chr11:121574272:TC>AA | N/A | p.F1123_R1124delinsLS | Inframe indel | 22.9 | | N/A | N/A | N/A | 72 | 409.453 |
| indels | chr11:chr11:121574272:TC>AA | N/A | p.F1123_R1124delinsLS | Inframe indel | 22.9 | | N/A | N/A | N/A | 117 | 370.733 |
| indels | chr11:chr11:121574272:TC>AA | N/A | p.F1123_R1124delinsLS | Inframe indel | 22.9 | | N/A | N/A | N/A | 108 | 299.198 |
| indels | chr11:chr11:121574272:TC>AA | N/A | p.F1123_R1124delinsLS | Inframe indel | 22.9 | | N/A | N/A | N/A | 33 | 295.017 |
| indels | chr11:chr11:121574272:TC>AA | N/A | p.F1123_R1124delinsLS | Inframe indel | 22.9 | | N/A | N/A | N/A | 55 | 263.975 |
| indels | chr11:121522971:T>TGGA | N/A | p.G527_A528insG | Inframe indel | 20.8 | | N/A | N/A | N/A | N/A | 330.769 |
| indels | chr11:121522971:T>TGGA | N/A | p.G527_A528insG | Inframe indel | 20.8 | | N/A | N/A | N/A | N/A | 174.635 |
| indels | chr11:121588028:T>TGTG | N/A | p.C1275_T1276insG | Inframe indel | 17.5 | | N/A | N/A | N/A | N/A | 377.034 |
| MPV | chr11:chr11:121570253:G>A | CR | p.S1107N | nonsynonymous SNV | 27.1 | | 0.775 | 1.17 | 1.69 | 87 | 257.705 |
| MPV | chr11:121543609:G>T | VPS10p | p.V583L | nonsynonymous SNV | 32 | | 0.476 | 1.17 | 1.69 | N/A | 338.905 |
| MPV | chr11:121574272:TC>AA | CR | p.F1123L | nonsynonymous SNV | 24.8 | | 0.859 | 1.17 | 1.69 | N/A | 424.498 |
| MPV | chr11:121577323:G>T | CR | p.G1168V | nonsynonymous SNV | 28.3 | | 0.782 | 1.17 | 1.69 | N/A | 361.02 |
| MPV | chr11:121605150:G>C | 3FN | p.W1563C | nonsynonymous SNV | 32 | | 0.476 | 1.17 | 1.69 | N/A | 450.547 |
| MPV | chr11:121574272:TC>AA &  chr11:121606902:T>C | CR & 3FN | p.F1123L & p.I1669T | Inframe indel & nonsynonymous SNV | 24.8 & 24.2 | 0.859 & 0.345 | | 1.17 | 1.69 | N/A | 145.168 |
| MPV | chr11:121608125:G>A | 3FN | p.G1730R | nonsynonymous SNV | 27.8 | | 0.673 | 1.17 | 1.69 | N/A | 207.044 |
| MPV | chr11:121625202:G>A | 3FN | p.V2097I | nonsynonymous SNV | 27.5 | | 0.209 | 1.17 | 1.69 | N/A | 540.573 |
| MPV | chr11:121625202:G>A | 3FN | p.V2097I | nonsynonymous SNV | 27.5 | | 0.209 | 1.17 | 1.69 | N/A | 311.246 |
| MPV | chr11:121625202:G>A | 3FN | p.V2097I | nonsynonymous SNV | 27.5 | | 0.209 | 1.17 | 1.69 | N/A | 534.211 |
| MPV | chr11:121627758:G>A | tail | p.D2190N | nonsynonymous SNV | 28.8 | | 0.42 | 1.17 | 1.69 | N/A | 491.498 |
| LPV | chr11:121554032:C>T | YWTD | p.R788W | nonsynonymous SNV | 27.8 | | 0.698 | 1.38 | 1.05 | 29 | 311.572 |
| LPV | chr11:121554032:C>T | YWTD | p.R788W | nonsynonymous SNV | 27.8 | | 0.698 | 1.38 | 1.05 | N/A | 220.277 |
| LPV | chr11:121558646:A>G | YWTD | p.M907V | nonsynonymous SNV | 24.3 | | 0.895 | 1.38 | 1.05 | N/A | 178.282 |
| LPV | chr11:121574261:G>T | CR | p.D1120Y | nonsynonymous SNV | 26.8 | | 0.64 | 1.38 | 1.05 | N/A | 386.986 |
| LPV | chr11:121589363:G>A | CR | p.G1351S | nonsynonymous SNV | 29.1 | | 0.614 | 1.38 | 1.05 | N/A | 409.438 |
| LPV | chr11:121621172:G>A | 3FN | p.G2000R | nonsynonymous SNV | 26.9 | | 0.555 | 1.38 | 1.05 | N/A | 484.537 |
| NPV | chr11:chr11:121478133:G>A | VPS10p | p.D140N | nonsynonymous SNV | 27.7 | | 0.404 | 1.15 | 1.09 | 120 | 391.672 |
| NPV | chr11:chr11:121514222:A>C | VPS10p | p.N371T | nonsynonymous SNV | 24.1 | | 0.297 | 1.15 | 1.09 | 63 | 333.412 |
| NPV | chr11:chr11:121543677:G>C | VPS10p | p.E605D | nonsynonymous SNV | 16.7 | | 0.057 | 1.15 | 1.09 | 74 | 239.198 |
| NPV | chr11:chr11:121543677:G>C | VPS10p | p.E605D | nonsynonymous SNV | 16.7 | | 0.057 | 1.15 | 1.09 | 171 | 410.451 |
| NPV | chr11:chr11:121545284:T>A | CC | p.S636T | nonsynonymous SNV | 24.9 | | 0.181 | 1.15 | 1.09 | 86 | 442.147 |
| NPV | chr11:chr11:121570213:A>G | CR | p.I1094V | nonsynonymous SNV | 13.4 | | 0.229 | 1.15 | 1.09 | 131 | 388.264 |
| NPV | chr11:chr11:121606865:C>T | 3FN | p.L1657F | nonsynonymous SNV | 33 | | 0.334 | 1.15 | 1.09 | 81 | 761.223 |
| NPV | chr11:chr11:121608123:G>A | 3FN | p.R1729H | nonsynonymous SNV | 31 | | 0.36 | 1.15 | 1.09 | N/A | 294.449 |
| NPV | chr11:chr11:121615047:C>T | 3FN | p.R1866W | nonsynonymous SNV | 34 | | 0.15 | 1.15 | 1.09 | N/A | 440.497 |
| NPV | chr11:121514164:G>A &  chr11:121520821:C>A | VPS10p | p.A352T & p.T459K | nonsynonymous SNV | 32 & … | | 0.398 & 0.118 | 1.15 | 1.09 | N/A | 202.966 |
| NPV | chr11:121550632:C>T | CC | p.A743V | nonsynonymous SNV | 20.2 | | 0.169 | 1.15 | 1.09 | N/A | 424.086 |
| NPV | chr11:121583575:T>G | CR | p.V1233G | nonsynonymous SNV | 16.1 | | 0.093 | 1.15 | 1.09 | N/A | 663.495 |
| NPV | chr11:121558767:C>T &  chr11:121618816:C>T &  chr11:121625161:A>G | YWTD & 3FN | p.T947M & p.R1883C & p.K2083R | nonsynonymous SNV | … & 22.3 & … | | 0.048 & 0.336 & 0.312 | 1.15 | 1.09 | N/A | 374.064 |
| NPV | chr11:121543677:G>C | VPS10p | p.E605D | nonsynonymous SNV | 16.7 | | 0.057 | 1.15 | 1.09 | N/A | 429.732 |
| p.E270K | chr11:121496918:G>A | VPS10p | p.E270K | nonsynonymous SNV | N/A | | 0.310 | 0.94 | 1.1 | N/A | 599.731 |
| p.E270K | chr11:121496918:G>A | VPS10p | p.E270K | nonsynonymous SNV | N/A | | 0.310 | 0.94 | 1.1 | N/A | 401.536 |
| p.E270K | chr11:121496918:G>A | VPS10p | p.E270K | nonsynonymous SNV | N/A | | 0.310 | 0.94 | 1.1 | N/A | 522.846 |
| p.E270K | chr11:121496918:G>A | VPS10p | p.E270K | nonsynonymous SNV | N/A | | 0.310 | 0.94 | 1.1 | N/A | 107.624 |
| p.E270K | chr11:121496918:G>A | VPS10p | p.E270K | nonsynonymous SNV | N/A | | 0.310 | 0.94 | 1.1 | N/A | 261.788 |
| p.E270K | chr11:121496918:G>A | VPS10p | p.E270K | nonsynonymous SNV | N/A | | 0.310 | 0.94 | 1.1 | N/A | 314.976 |
| p.E270K | chr11:121496918:G>A | VPS10p | p.E270K | nonsynonymous SNV | N/A | | 0.310 | 0.94 | 1.1 | N/A | 590.453 |
| p.E270K | chr11:121496918:G>A | VPS10p | p.E270K | nonsynonymous SNV | N/A | | 0.310 | 0.94 | 1.1 | N/A | 401.034 |
| p.E270K | chr11:121496918:G>A | VPS10p | p.E270K | nonsynonymous SNV | N/A | | 0.310 | 0.94 | 1.1 | N/A | 376.459 |
| p.E270K | chr11:121496918:G>A | VPS10p | p.E270K | nonsynonymous SNV | N/A | | 0.310 | 0.94 | 1.1 | N/A | 487.447 |
| p.E270K | chr11:121496918:G>A | VPS10p | p.E270K | nonsynonymous SNV | N/A | | 0.310 | 0.94 | 1.1 | N/A | 481.864 |
| p.A528T | chr11:121522975:G>A | VPS10p | p.A528T | nonsynonymous SNV | N/A | | 0.112 | 1.1 | 1.2 | 136 | 681.917 |
| p.A528T | chr11:121522975:G>A | VPS10p | p.A528T | nonsynonymous SNV | N/A | | 0.112 | 1.1 | 1.2 | N/A | 339.444 |
| p.A528T | chr11:121522975:G>A | VPS10p | p.A528T | nonsynonymous SNV | N/A | | 0.112 | 1.1 | 1.2 | N/A | 378.698 |
| p.A528T | chr11:121522975:G>A | VPS10p | p.A528T | nonsynonymous SNV | N/A | | 0.112 | 1.1 | 1.2 | N/A | 270.399 |
| p.A528T | chr11:121522975:G>A | VPS10p | p.A528T | nonsynonymous SNV | N/A | | 0.112 | 1.1 | 1.2 | N/A | 481.567 |
| p.A528T | chr11:121522975:G>A | VPS10p | p.A528T | nonsynonymous SNV | N/A | | 0.112 | 1.1 | 1.2 | N/A | 361.7 |
| p.A528T | chr11:121522975:G>A | VPS10p | p.A528T | nonsynonymous SNV | N/A | | 0.112 | 1.1 | 1.2 | N/A | 441.828 |
| p.A528T | chr11:121522975:G>A | VPS10p | p.A528T | nonsynonymous SNV | N/A | | 0.112 | 1.1 | 1.2 | N/A | 652.189 |
| p.A528T | chr11:121522975:G>A | VPS10p | p.A528T | nonsynonymous SNV | N/A | | 0.112 | 1.1 | 1.2 | N/A | 450.962 |
| p.A528T | chr11:121522975:G>A | VPS10p | p.A528T | nonsynonymous SNV | N/A | | 0.112 | 1.1 | 1.2 | N/A | 575.171 |
| p.A528T | chr11:121522975:G>A | VPS10p | p.A528T | nonsynonymous SNV | N/A | | 0.112 | 1.1 | 1.2 | N/A | 481.263 |
| p.A528T + p.E270K | chr11:121496918:G>A &  chr11:121522975:G>A | VPS10p | p.E270K & p.A528T | nonsynonymous SNV | N/A | | 0.310 & 0.112 | N/A | N/A | N/A | 443.738 |
| p.A528T + p.E270K | chr11:121496918:G>A &  chr11:121522975:G>A | VPS10p | p.E270K & p.A528T | nonsynonymous SNV | N/A | | 0.310 & 0.112 | N/A | N/A | N/A | 470.032 |

P. cons = protein consequence, EOAD = Early Onset Alzheimer’s Disease , LOAD = Late Onset Alzheimer’s Disease, WB rel levels = Western Blot relative levels.

The E270K and A528T variants have previously been associated with increased risk for LOAD (11).

### Supplemental Table 2. CSF-sSORL1 per pathogenicity category

| **comparison** | **Mean carriers** | **SD carriers** | **N cases** | **Mean contr.** | **SD contr.** | **N contr.** | **p** | **Effect (β)** | **SE** |
| --- | --- | --- | --- | --- | --- | --- | --- | --- | --- |
| PTV vs WT | 260 | 123 | 9 | 466 | 133 | 128 | 2.1E-07 | 237.4 | 45.8 |
| PTV vs AD WT | 260 | 123 | 9 | 462 | 134 | 78 | 5.3E-06 | 228.3 | 50.2 |
| PTV vs Contr | 260 | 123 | 9 | 474 | 131 | 50 | 5.2E-10 | 267.4 | 43.0 |
| HPV vs WT | 323 | 154 | 18 | 466 | 133 | 128 | 9.8E-08 | 171.8 | 32.2 |
| HPV vs AD WT | 323 | 154 | 18 | 462 | 134 | 78 | 2.1E-06 | 168.5 | 35.5 |
| HPV vs Contr | 323 | 154 | 18 | 474 | 131 | 50 | 7.9E-10 | 192.3 | 31.3 |
| MPV vs WT | 369 | 132 | 11 | 466 | 133 | 128 | 1.9E-02 | 94.4 | 40.3 |
| MPV vs AD WT | 369 | 132 | 11 | 462 | 134 | 78 | 4.0E-02 | 89.5 | 43.6 |
| MPV vs Contr | 369 | 132 | 11 | 474 | 131 | 50 | 1.1E-03 | 123.8 | 37.9 |
| LPV vs WT | 332 | 117 | 6 | 466 | 133 | 128 | 1.1E-02 | 134.7 | 53.1 |
| LPV vs AD WT | 332 | 117 | 6 | 462 | 134 | 78 | 1.4E-02 | 134.3 | 54.7 |
| LPV vs Contr | 332 | 117 | 6 | 474 | 131 | 50 | 1.2E-03 | 154.6 | 47.8 |
| NPV vs WT | 414 | 148 | 14 | 466 | 133 | 128 | 1.6E-01 | 51.2 | 36.6 |
| NPV vs AD WT | 414 | 148 | 14 | 462 | 134 | 78 | 1.6E-01 | 54.3 | 39.1 |
| NPV vs Contr | 414 | 148 | 14 | 474 | 131 | 50 | 7.1E-02 | 62.2 | 34.4 |
| p.A528T vs WT | 464 | 118 | 13 | 466 | 133 | 128 | 7.2E-01 | 13.5 | 37.3 |
| p.A528T vs AD WT | 464 | 118 | 13 | 462 | 134 | 78 | 8.8E-01 | 6.2 | 41.6 |
| p.A528T vs Contr | 464 | 118 | 13 | 474 | 131 | 50 | 1.4E-01 | 52.2 | 35.4 |
| p.E270K vs WT | 420 | 135 | 13 | 466 | 133 | 128 | 2.9E-01 | 40.3 | 38.4 |
| p.E270K vs AD WT | 420 | 135 | 13 | 462 | 134 | 78 | 3.8E-01 | 36.2 | 41.1 |
| p.E270K vs Contr | 420 | 135 | 13 | 474 | 131 | 50 | 1.1E-01 | 62.0 | 38.4 |
| Indels vs WT | 315 | 75 | 8 | 466 | 133 | 128 | 1.1E-03 | 152.5 | 46.8 |
| Indels vs AD WT | 315 | 75 | 8 | 462 | 134 | 78 | 2.4E-03 | 153.1 | 50.4 |
| Indels vs Contr | 315 | 75 | 8 | 474 | 131 | 50 | 1.9E-05 | 168.2 | 39.3 |
| AD WT vs Contr | 462 | 134 | 78 | 474 | 131 | 50 | 4.5E-01 | 18.9 | 25.1 |

Mean CSF-sSORL1 concentrations per pathogenicity category and statistics of robust linear regression of CSF-sSORL1 concentrations versus *SORL1*-WT (*SORL1*-WT AD + controls), *SORL1-*WT AD or *SORL1*-WT controls. SD = standard deviation , SE = Standard Error. PTV: protein truncating variants; HPV: high priority missense variants; MPV: moderate priority missense variants; LPV: low priority missense variants; NPV: no priority missense variants; AD WT: AD cases with wild type *SORL1*.

Supplemental figures

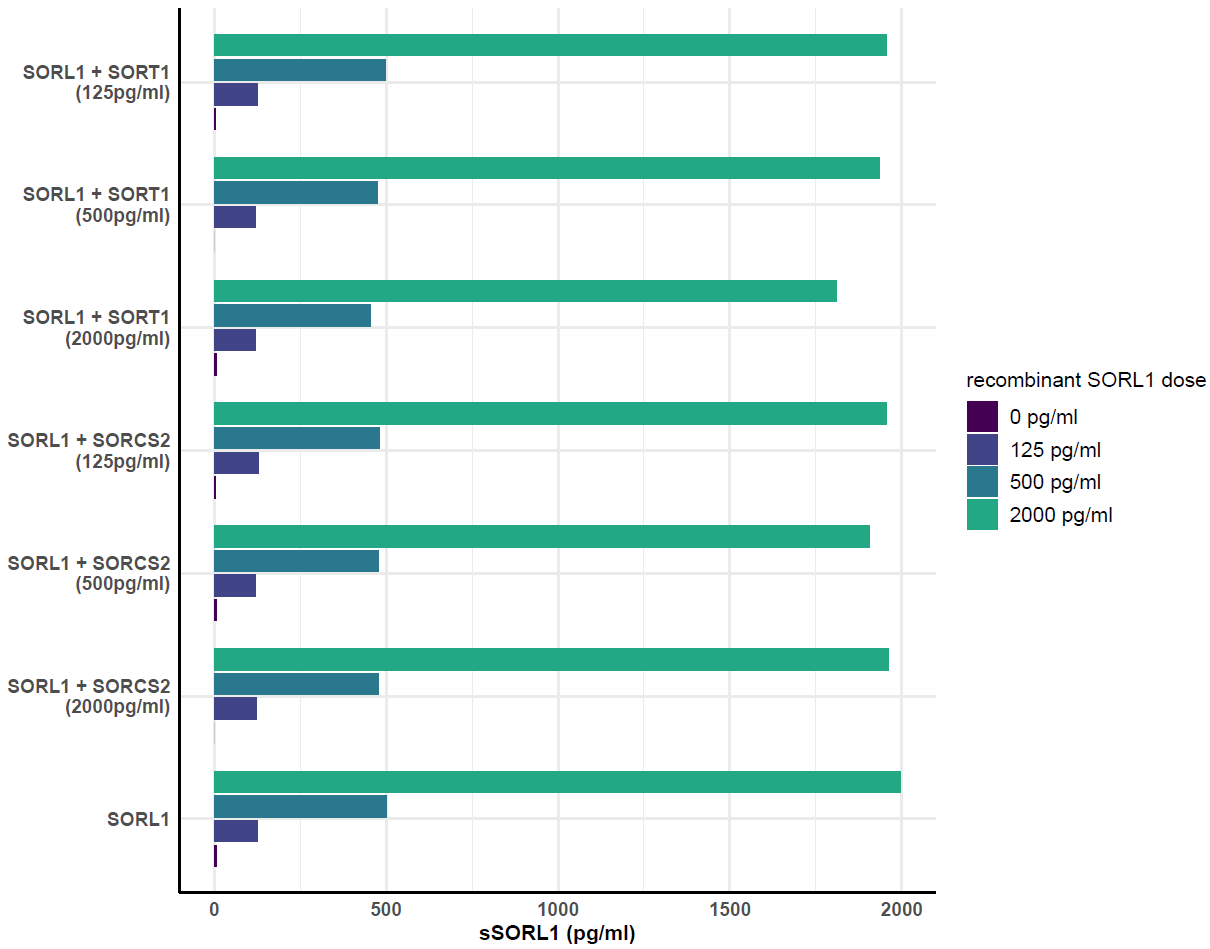

### Supplemental figure 1. Specificity/cross-reactivity across VPS10p-domain family members

Measurement of spiked-in levels of VPS10p-domain family proteins SORT1 and SORCS2. We tested four concentrations of recombinant SORL1 (blank, sample with 125 pg/ml, 500 pg/ml, and 2000 pg/ml) were spiked with varying concentrations of recombinant SORT1 or SORCS2 (125 pg/ml, 500 pg/ml, 2000 pg/ml).

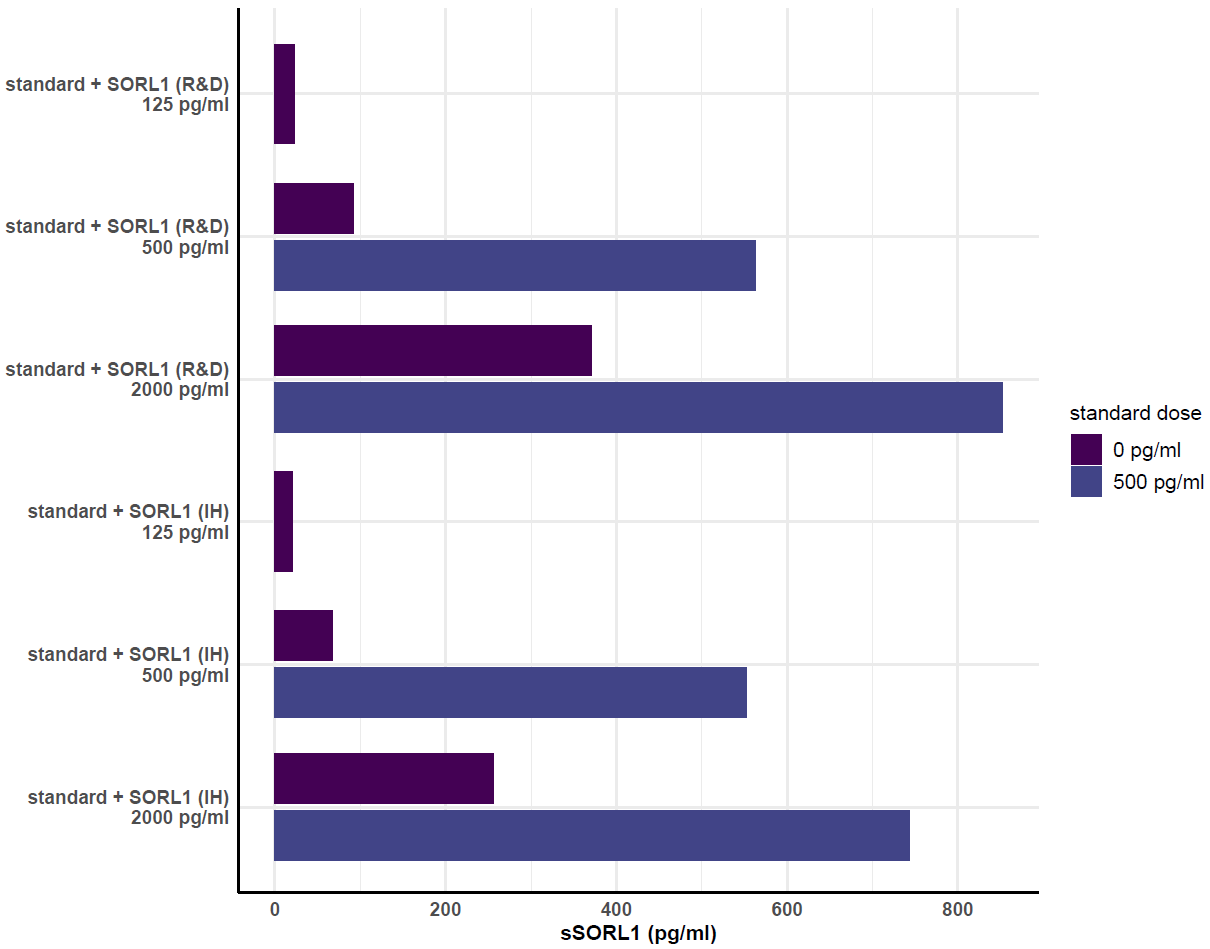

### Supplemental figure 2. Recognition of full recombinant SORL1 constructs

A blank sample was spiked with three different concentrations (125 pg/ml, 500 pg/ml, and 2000 pg/ml) of various full-length recombinant SORL1 proteins. Additionally, a sample containing 500 pg/ml of the standard recombinant from the kit was spiked with either 500 or 2000 pg/ml of full-length recombinant SORL1. SORL1 (IH) refers to the in-house produced SORL1 recombinant, while SORL1 (R&D) corresponds to the commercially available recombinant SORL1 from R&D Systems.

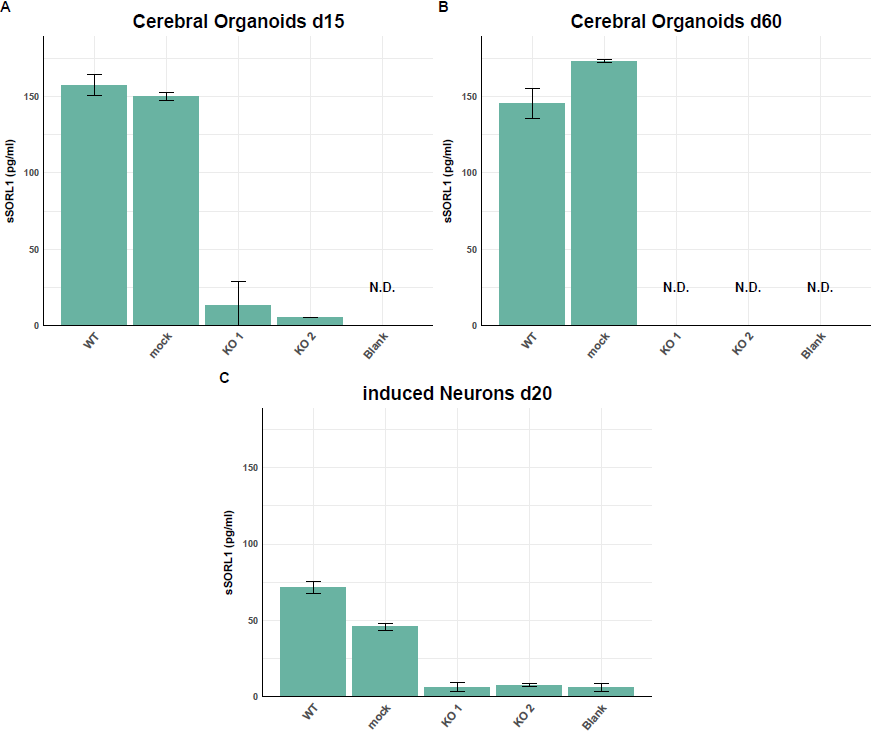

### Supplemental figure 3. sSORL1 in cell culture medium of SORL1 KO cells

ELISA measurements of sSORL1 in cell culture medium from various cell types, including 20-day-old induced neurons, 15-day-old cerebral organoids, and 60-day-old cerebral organoids. Each cell type included Wild Type, mock, and KO conditions. Mock cells served as positive controls, undergoing the full CRISPR workflow without any genetic modification. Overall, sSORL1 levels were lower in the induced neurons compared to the organoids.

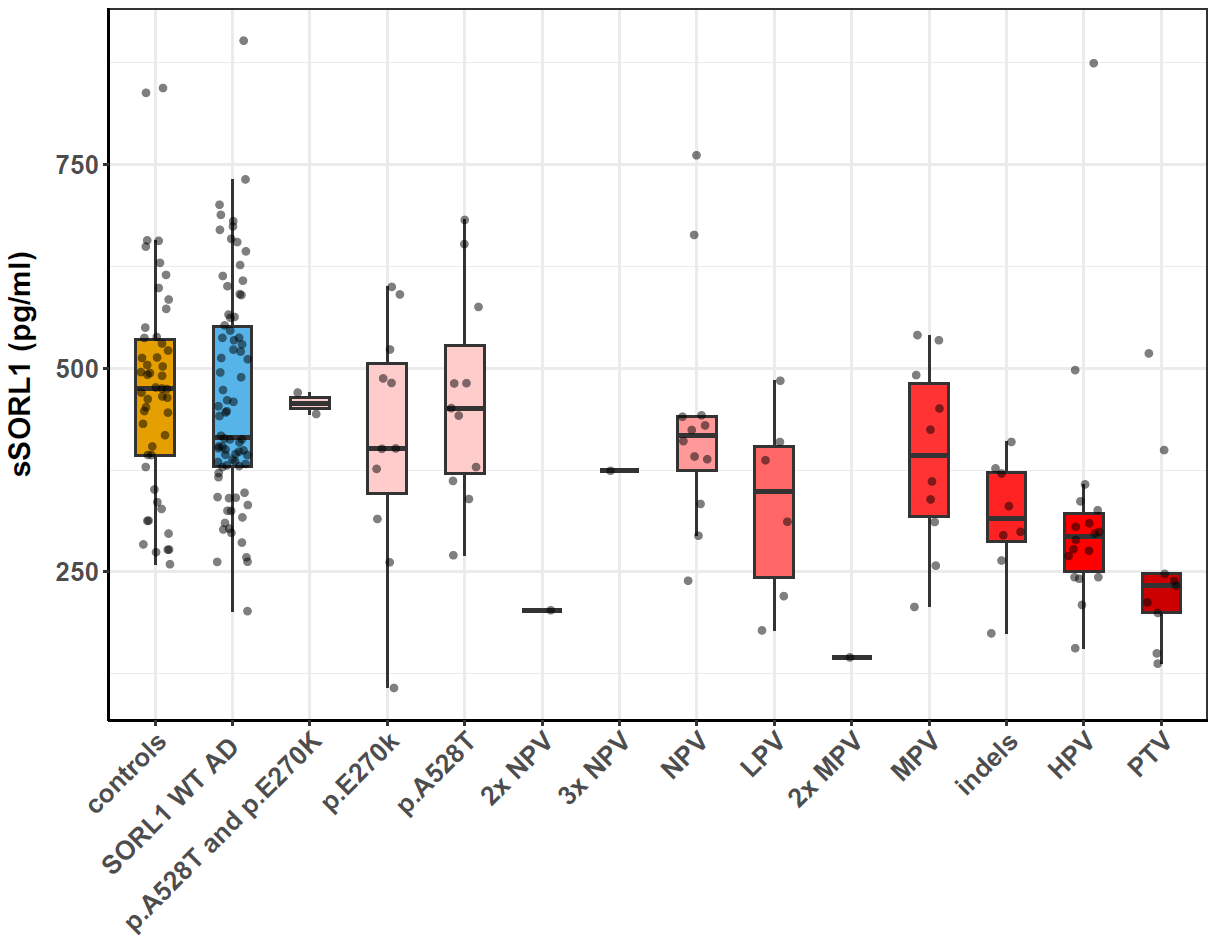

### Supplemental figure 4. CSF-sSORL1 in carriers of diverse *SORL1* variants

ELISA CSF measurements of the different *SORL1* variant carriers. Several AD cases carries more than one variant, which were represented in different categories here. Controls (N=50), SORL1 WT AD (N=78), p.A528T and p.E270K (N=2), p.E270K (N=11), p.A528T (N=11), 2x NPV (N=1), 3x NPV (N=1), NPV (N=12), LPV (N=6), 2x MPV (N=1), MPV (N=10), indels (N=8), HPV (N=18), PTV (N=9).

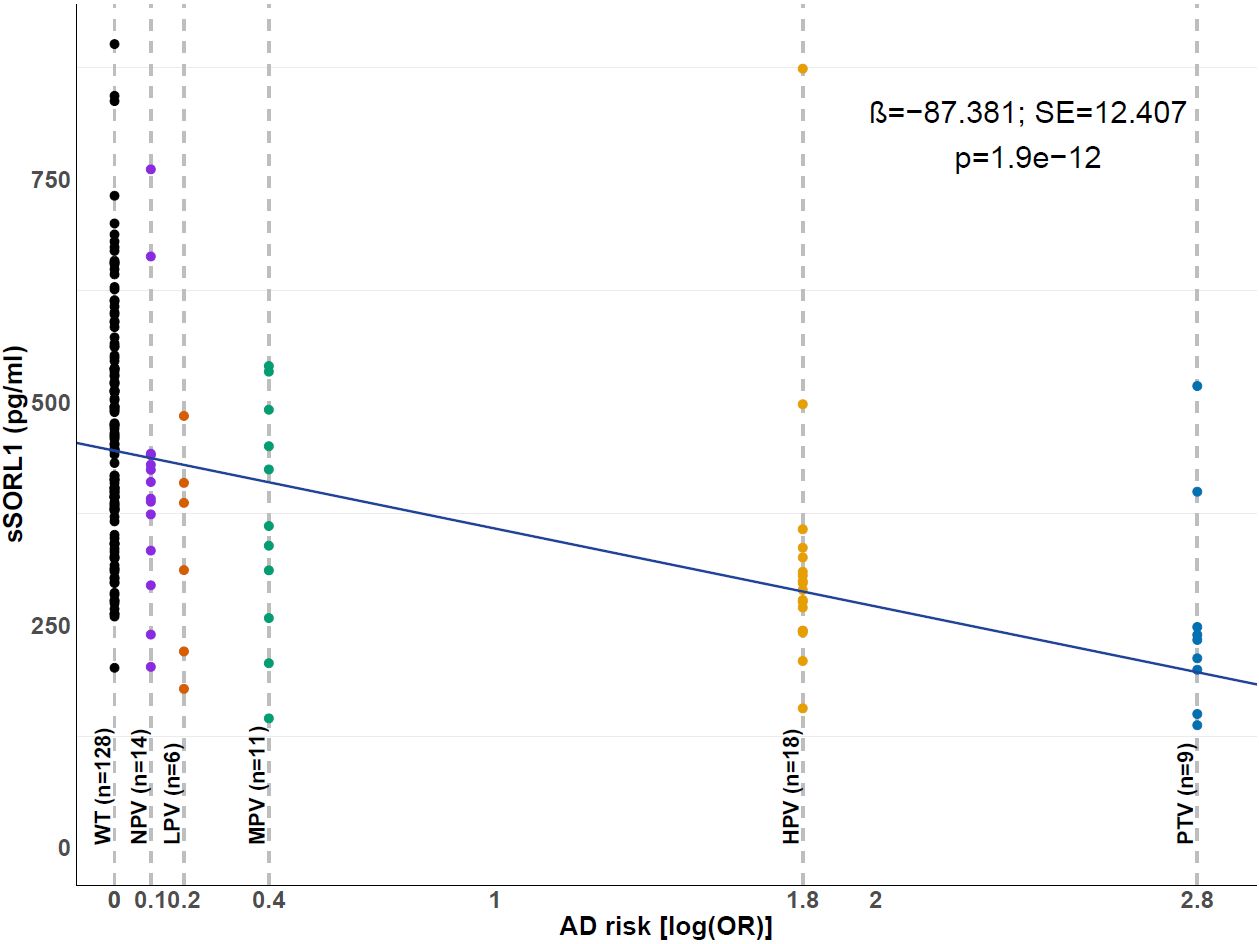

### Supplemental figure 5. Per SORL1 variant category: CSF-sSORL1 concentrations vs. AD risk

CSF-sSORL1 concentrations from carriers of high-risk variants are lower. Robust linear regression analysis comparing CSF-sSORL1 concentrations with the AD risk (OR) associated with each *SORL1* pathogenic category as reported in Holstege et al., 2024. The ORs for each category were as follows: PTV = 17.2, HPV = 6.1, MPV = 1.5, LPV = 1.2, and NPV = 1.1, SORL1 WT controls = 1. The natural logarithm of these ORs was used on the x-axis of the analysis (i.e. PTV= 2.8, HPV= 1.8, MPV = 0.4, LPV = 0.2, NPV = 0.1, WT = 0).

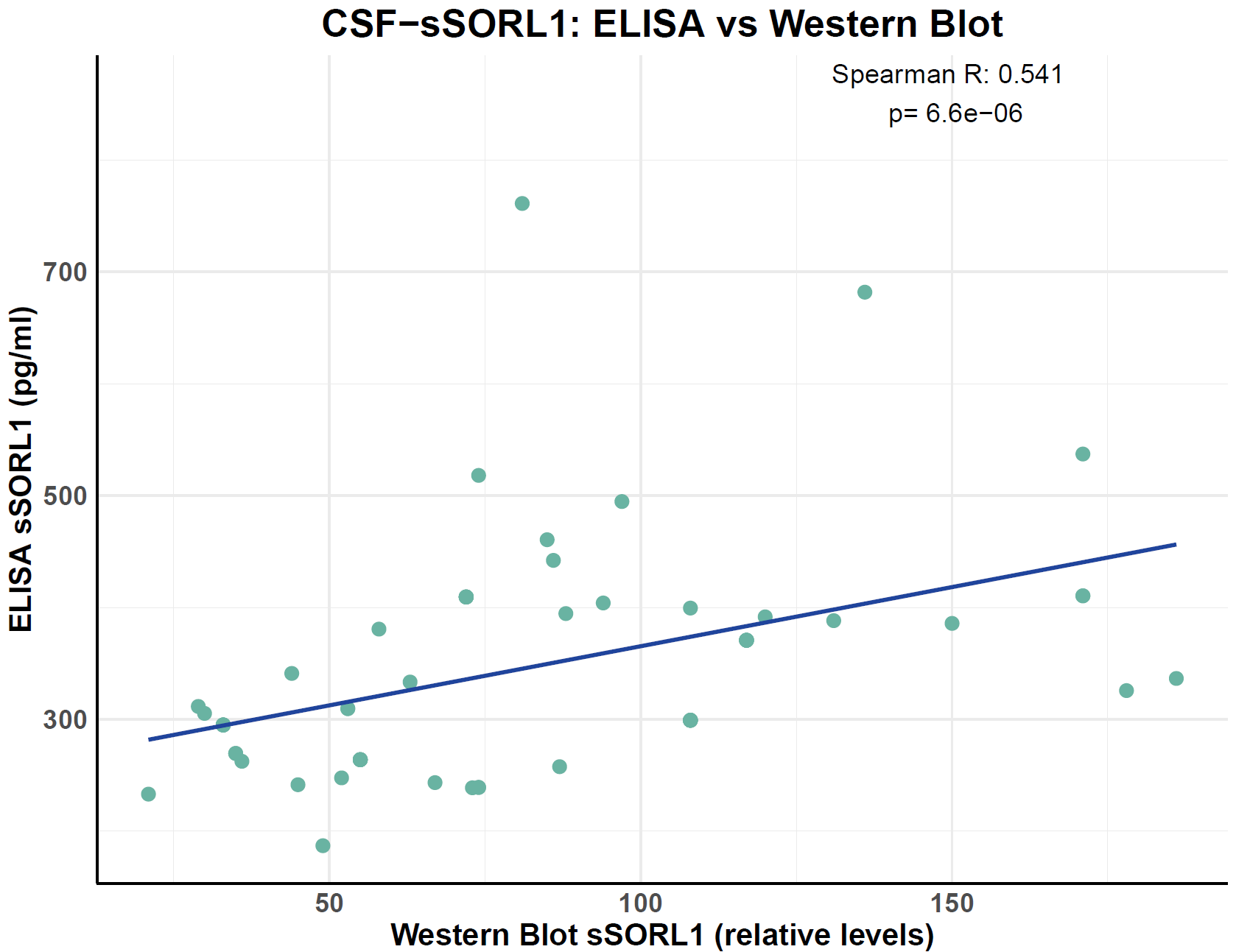

### Supplemental figure 6. CSF-sSORL1: ELISA vs Western Blot

Spearman correlations between ELISA-derived sSORL1 levels and Western blot-based relative sSORL1 levels. The left panel shows the correlation of absolute ELISA sSORL1 levels, while the right panel displays the correlation of relative ELISA sSORL1 levels.

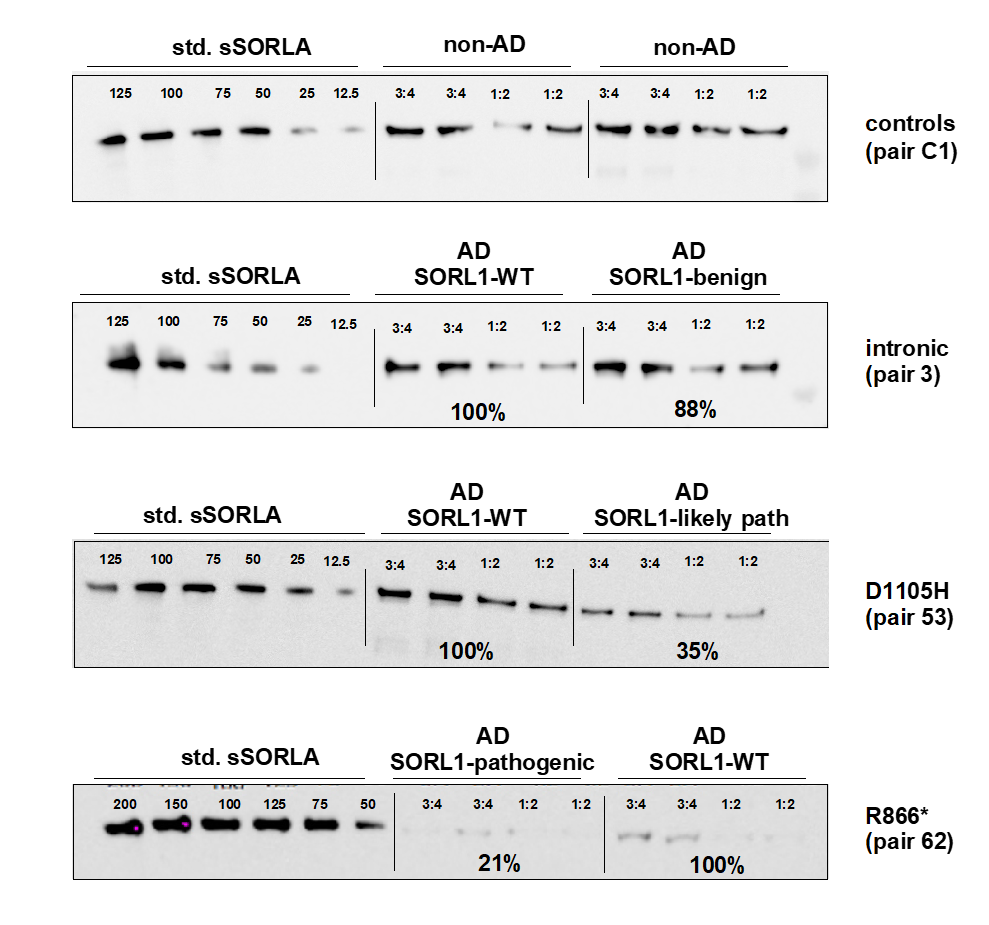

### Supplemental figure 7. CSF-sSORL1 in AD patients with/without SORL1 genetic variants

Representative Western blots of CSF samples from AD patients who carry *SORL1* genetic variants from different pathogenicity groups and their uniquely matched *SORL1*-WT AD control. A ~250 kDa band corresponding to sSORL1 was observed across different carriers. Samples to the left of each blot correspond to internal standards of recombinant purified sSORL1 protein at variable concentrations (between 125 and 12.5 ng/mL). The CSF samples from Pair C1 originates from two non-AD individuals, Pair 3 (‘benign’ *SORL1* intronic variant 121.393.721 G>A), Pair 53 (high priority or ‘likely pathogenic’ *SORL1* variant 121.440.955 G>C; p.D1105H) and Pair 62 (‘pathogenic’ truncating *SORL1* variant R866*).

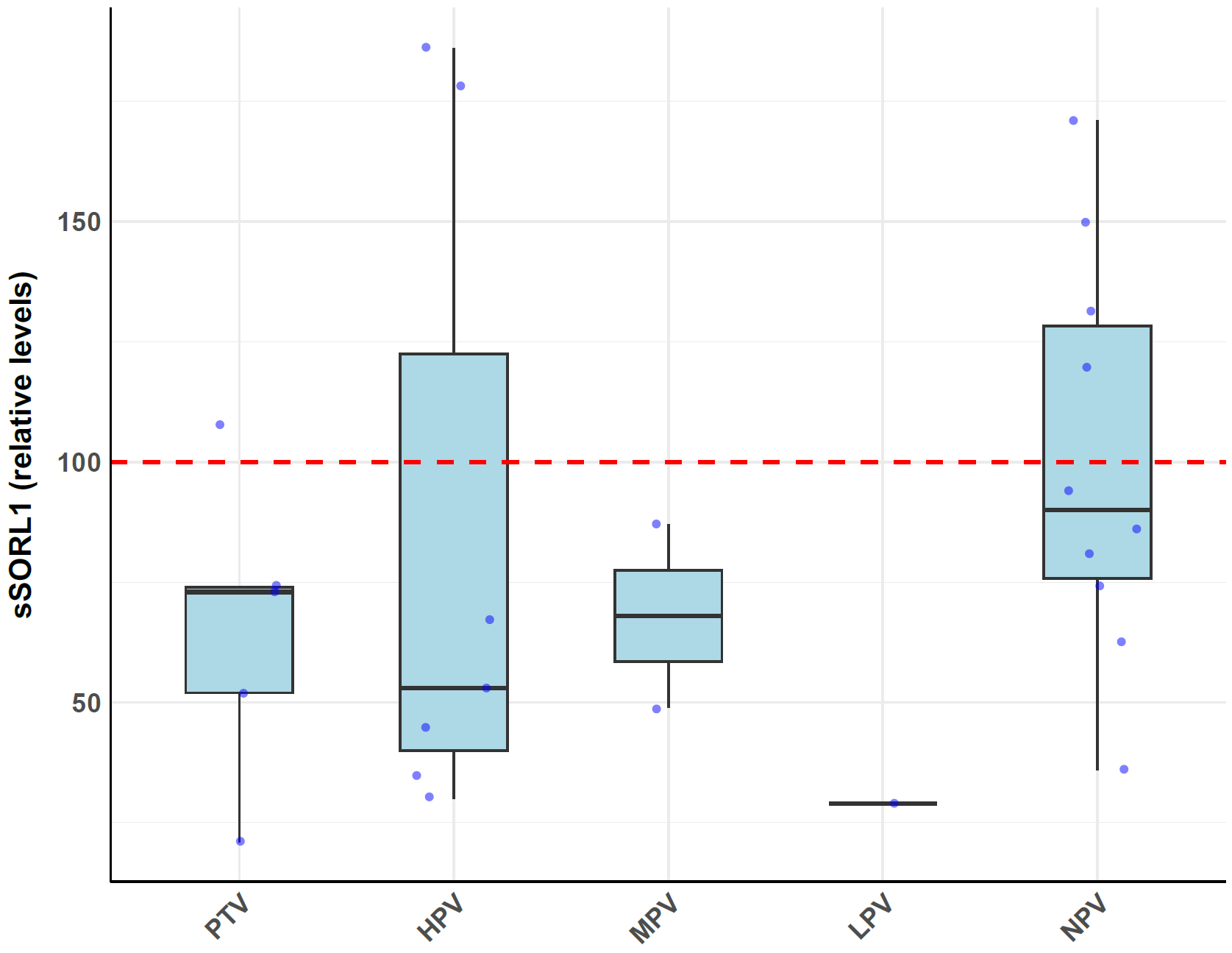

### Supplemental figure 8. Western Blot: CSF-sSORL1 per variant priority group

Western blotting was performed to evaluate the relative expression of proteins in the priority groups. The red dotted line indicates the baseline control level for comparison. Relative expression levels for each sample were determined by normalizing the protein signal of each sample to its corresponding control. While no pathogenicity group showed a statistically significant decrease compared to the reference median, most pathogenic variant carriers exhibited lower expression levels than their matched controls.

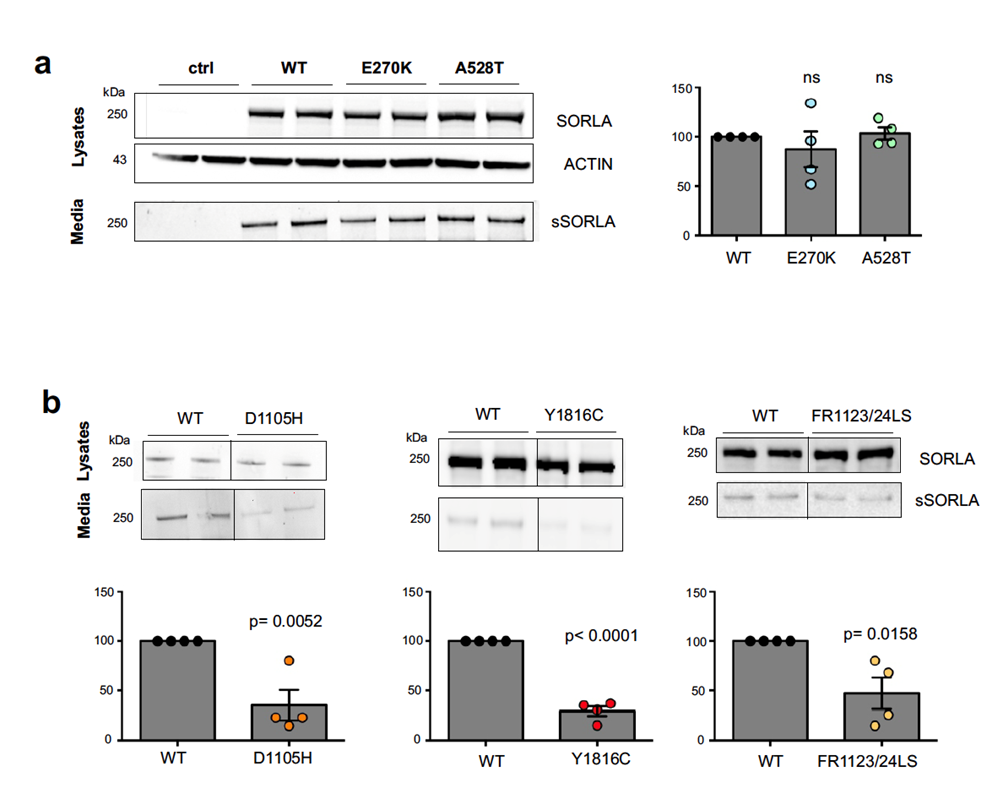

### Supplemental figure 9. Western Blot: CSF-sSORL1 in HPV vs common variants carriers

Representative WB of lysates and media samples from N2a cells transfected with *SORL1*-wild type (WT) or *SORL1*-variants p.E270K and p.A528K (a), p.D1105H (b), p.Y1816C (c), or p.FR1123/24LS (d). The amount of shed sSORL1 ectodomain was quantified by densiometric scanning of the blots and the signals for the mutant construct were expressed as relative to the WT signal. The means of four independent experiments for each variant are shown ± SEM. Comparisons of sSORL1 concentrations between mutated and wild-type (WT) *SORL1* conditions were performed using a two-tailed Student’s t-test.

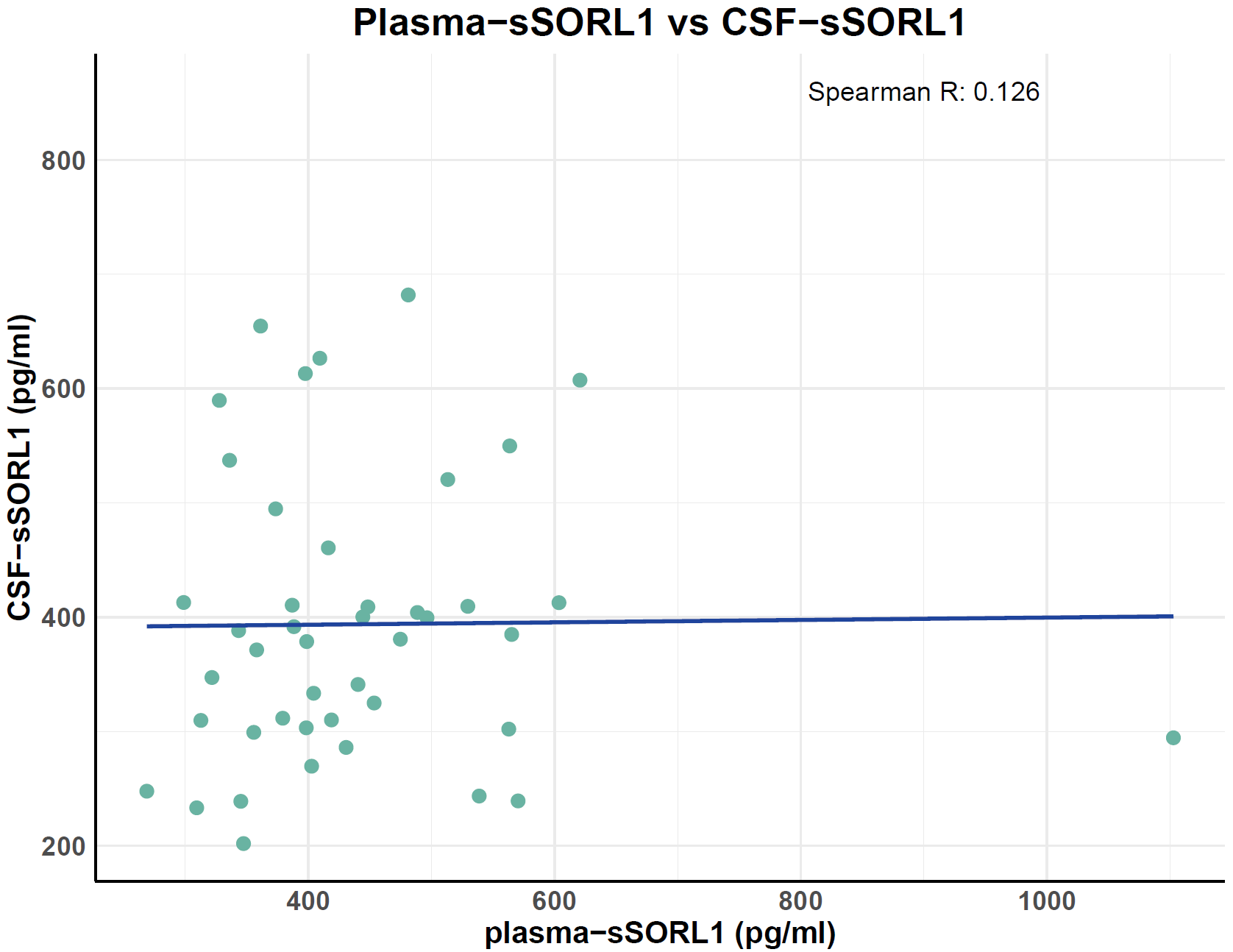

### Supplemental figure 10. Plasma-sSORL1 vs CSF-sSORL1 concentrations

Spearman correlation between CSF sSORL1 concentrations and plasma sSORL1 concentrations in 44 individuals.

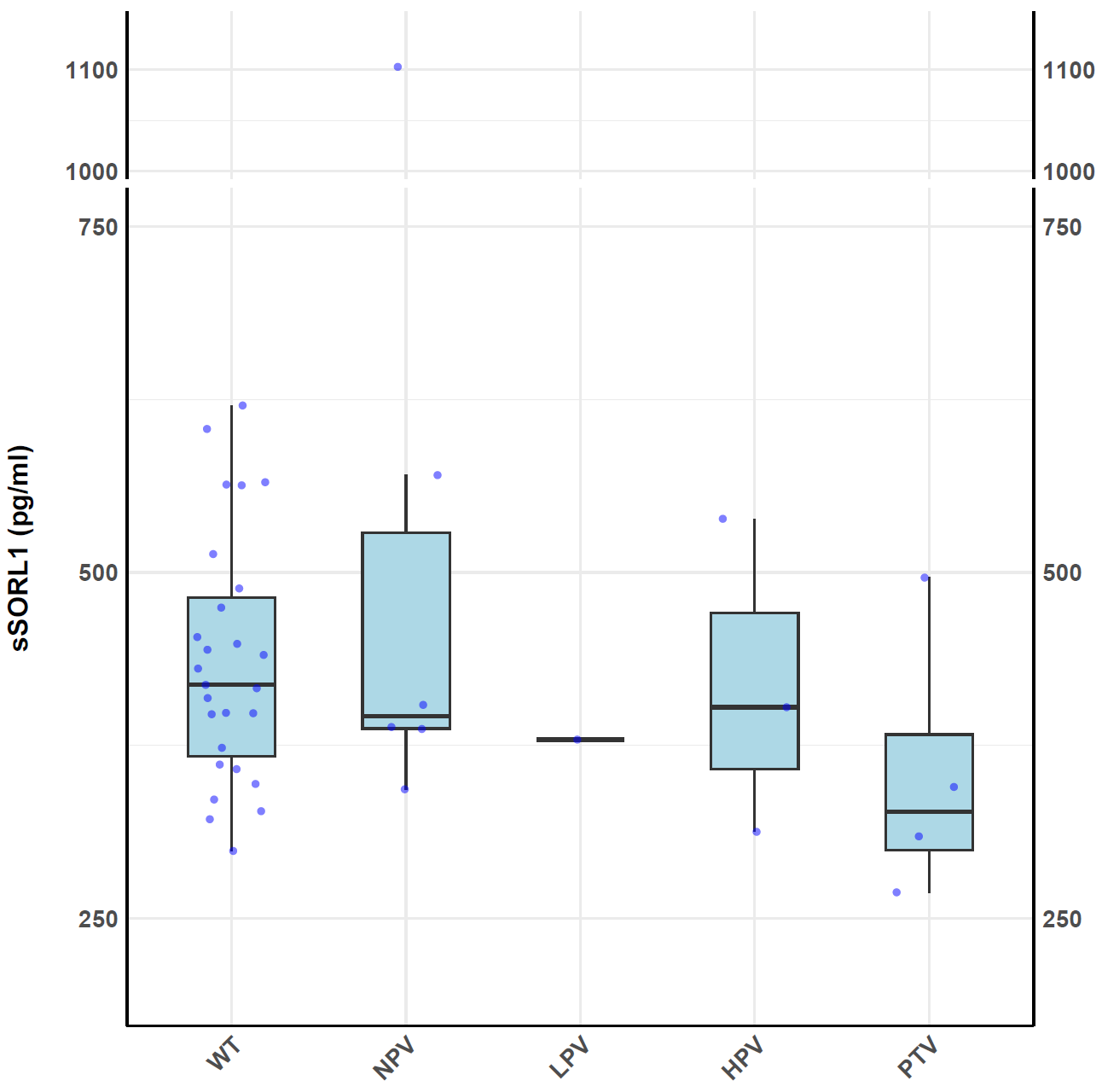

### Supplemental figure 11. Plasma-sSORL1 concentrations per variant group

ELISA Plasma-sSORL1 measurements of the different *SORL1* variant carriers.

Individuals consisted of 27 non-carriers, 6 NPV, 1 LPV, 3 HPV, 4 PTV. No significant difference was found between pathogenicity categories.

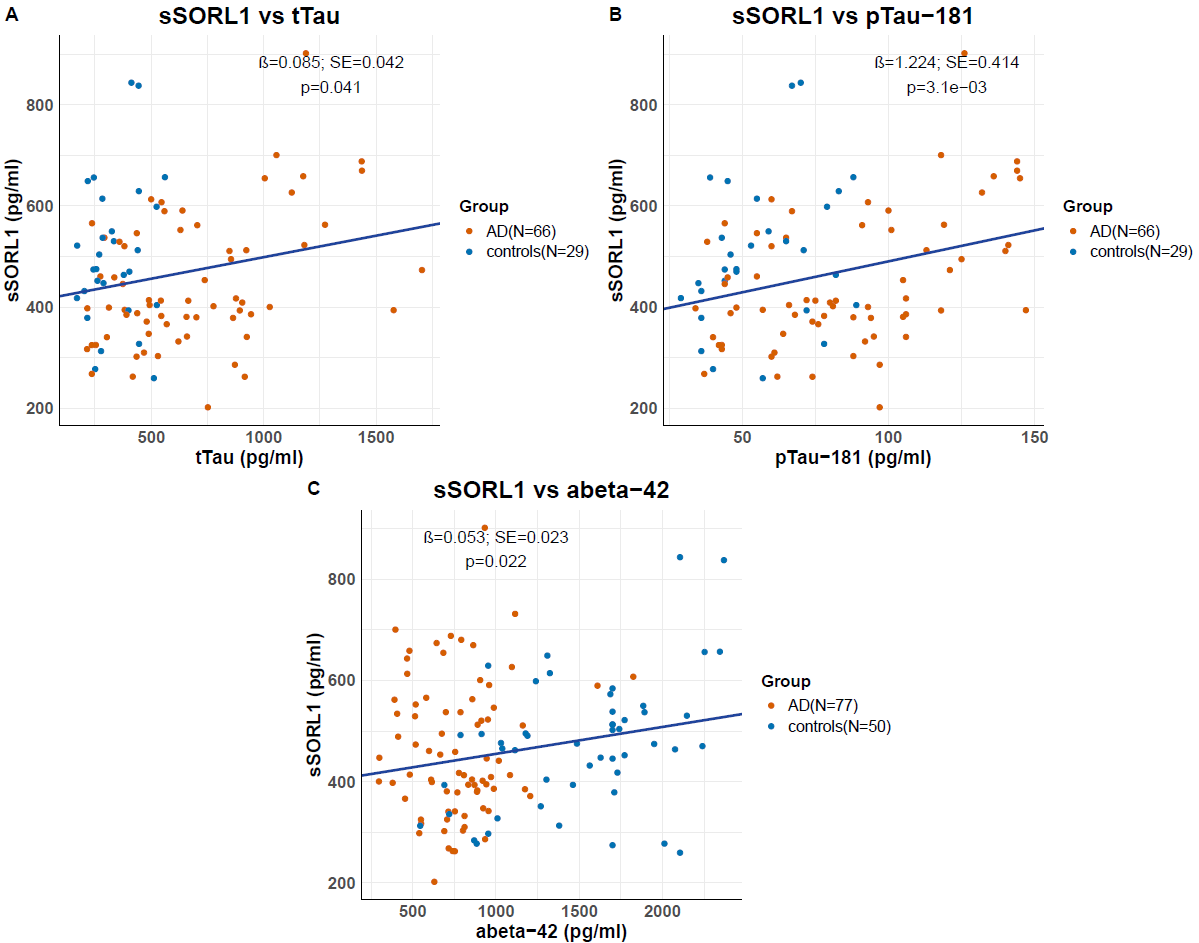

### Supplemental figure 12. CSF-sSORL1 vs AD CSF-biomarkers in context of WT *SORL1*

To assess the relationship between CSF-sSORL1 and tTau (n=95) (A), pTau-181 (n=95) (B) and Aβ42 (n=127) (C), we performed a Robust Linear Model to account for the potential influence of outliers across AD cases and controls. Fewer observations were available for tTau and pTau-181 compared to Aβ42. The association of the AD biomarkers with sSORL1 were corrected for age of collection and sex.
